# Transcriptional regulation of the *N*_ε_-fructoselysine metabolism in *Escherichia coli* by global and substrate-specific cues

**DOI:** 10.1101/2020.01.13.904318

**Authors:** Benedikt Graf von Armansperg, Franziska Koller, Nicola Gericke, Michael Hellwig, Pravin Kumar Ankush Jagtap, Ralf Heermann, Janosch Hennig, Thomas Henle, Jürgen Lassak

## Abstract

Thermally processed food is an important part of the human diet. Heat-treatment, however, promotes the formation of so-called Amadori rearrangement products (ARPs), such as fructoselysine. The gut microbiota including *Escherichia coli* can utilize these compounds as a nutrient source. While the degradation route for fructoselysine is well described, regulation of the corresponding pathway genes *frlABCD* remained poorly understood. Here we use bioinformatics combined with molecular and biochemical analyses and show that in *E. coli*, fructoselysine metabolism is tightly controlled at the transcriptional level. The global regulator Crp (CAP), as well as the alternative sigma factor σ32 (RpoH) contribute to promoter activation at high cAMP-levels and heat stress, respectively. In addition, we identified and characterized a transcriptional regulator FrlR, encoded adjacent to *frlABCD*, as fructoselysine-6-phosphate specific roadblock repressor. Our study provides profound evidence that the interplay of global and substrate-specific regulation is a perfect adaptation strategy to efficiently utilize unusual substrates within the human gut environment.

**Abbreviated Summary:** Thermal food processing promotes the formation of Amadori rearrangement products (ARPs), such as fructoselysine. The gut microbiota including *Escherichia coli* can utilize these compounds as a nutrient source. We show that in *E. coli*, fructoselysine metabolism is tightly controlled at the transcriptional level by global and substrate-specific regulators. Their interplay is a perfect adaptation strategy to efficiently utilize fructoselysine within the human gut environment.

**Graphical Abstract:** 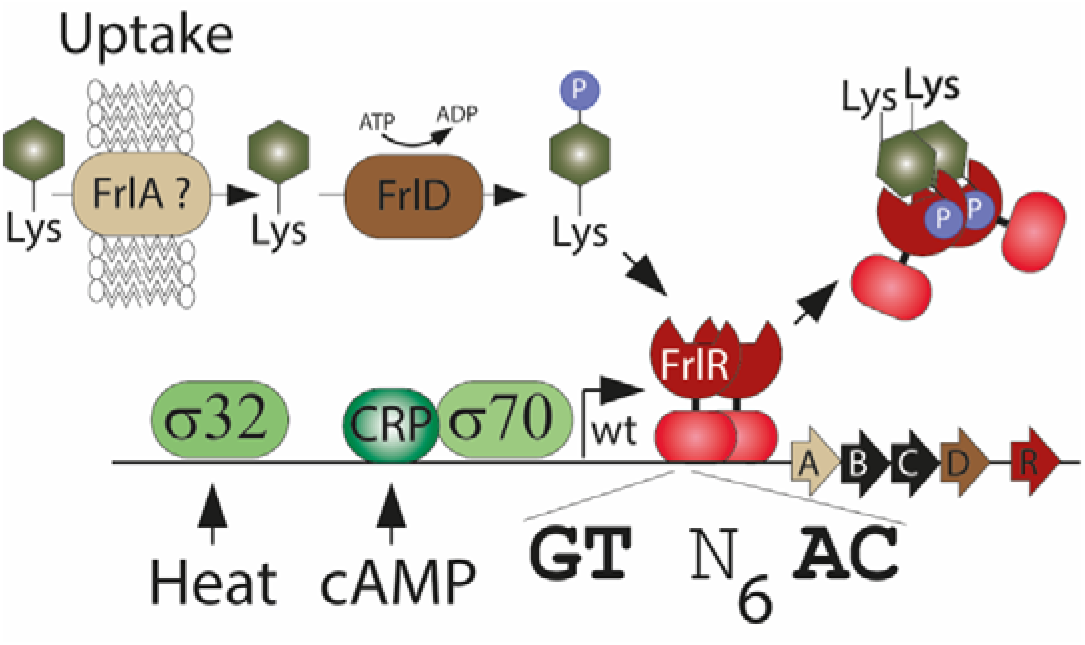

## Introduction

Glycation is a non-enzymatic form of glycosylation and means a spontaneous reaction of amino compounds with reducing sugars such as glucose (Ulrich & Cerami, 2001, Lassak *et al.*, 2019). The phenomenon was first described by Louis-Camille Maillard in 1912 being predominantly responsible for the taste, aroma and appearance of thermally processed food (Maillard, 1912a, Maillard, 1912b). Simple condensation products of primary amino groups at the N-termini of polypeptides or the e-amino groups of lysine and reducing sugars such as glucose are the most prevalent Maillard reaction products in food (Henle, 2003). These “sugar-amino acids” are also called “Amadori rearrangement products” (ARPs) (Amadori, 1925, Hodge, 1955). Protein-bound Maillard reaction products can be the result of a condensation reaction, taking place between an aldose or ketose and a primary amine either in form of an α-amino group at the N-terminus or an ε-amino group of lysine residues within the polypeptide chain. Bacteria, including *Escherichia coli*, *Bacillus subtilis* and *Salmonella enterica* have evolved efficient strategies to use these ARPs as sole carbon source (Wiame *et al.*, 2002, Wiame *et al.*, 2004, Wiame *et al.*, 2005, Ali *et al.*, 2014, Miller *et al.*, 2015). In this regard, utilization specifically of fructose asparagine by the food borne pathogen *S. enterica* provides a fitness advantage in the inflamed intestine (Ali *et al.*, 2014). Notably, the wide distribution of the ARP catabolism among the gut microbiota (Sabag-Daigle *et al.*, 2018) further suggests, that this carbon source might play an important role in colonizing the intestinal environment.

While there are numerous uptake mechanisms and diverse specificity towards glycation products in distinct microorganisms (Wiame *et al.*, 2004, Miller *et al.*, 2015, Sabag-Daigle *et al.*, 2017), the degradation route follows a conserved route and can be illustrated by the *E. coli N*_ε_*-*fructoselysine (ε-FrK) metabolism. Upon uptake - presumably by the putative permease FrlA - a kinase FrlD phosphorylates the sugar moiety at the C6-position (Fig. 1) (Wiame *et al.*, 2002). In a second step, the deglycase FrlB hydrolyses fructoselysine-6-phosphate into glucose-6-phosphate and lysine to be further processed via glycolysis and amino acid metabolism, respectively. Together with an additional *N*_ε_-psicoselysine/ε-FrK epimerase FrlC (Wiame & Van Schaftingen, 2004), the pathway is encoded in one single operon *frlABCD* of thus far unknown regulation. In the present study we show, that the *E. coli* FrK catabolism is tightly controlled by positive and negative regulation. On the one hand, the global transcription factor Crp (CAP) as well as the sigma factor σ^32^ (RpoH) contribute to promoter activation at high cAMP-levels and heat stress, respectively. On the other hand, we identified the previously elusive regulator FrlR, encoded adjacent to *frlABCD*, as ε-FrK specific roadblock repressor. However, ε-FrK is not recognized directly but only upon phosphorylation. FrlR itself is a member of the GntR/HutC family of transcriptional regulators recognizing the consensus sequence GT(N)_x_AC. Binding of ε-FrK-leads to structural rearrangements in the FrlR transcriptionally active dimer, which in turn weakens DNA binding, and subsequently permits transcription of the *frlABCD* operon. Thus, we conclude that the interplay of global and substrate-specific regulation combined with a heat stress dependent transcriptional upshift is a perfect adaptation to utilize thermally processed food within the human gut environment.

**Fig 1:**
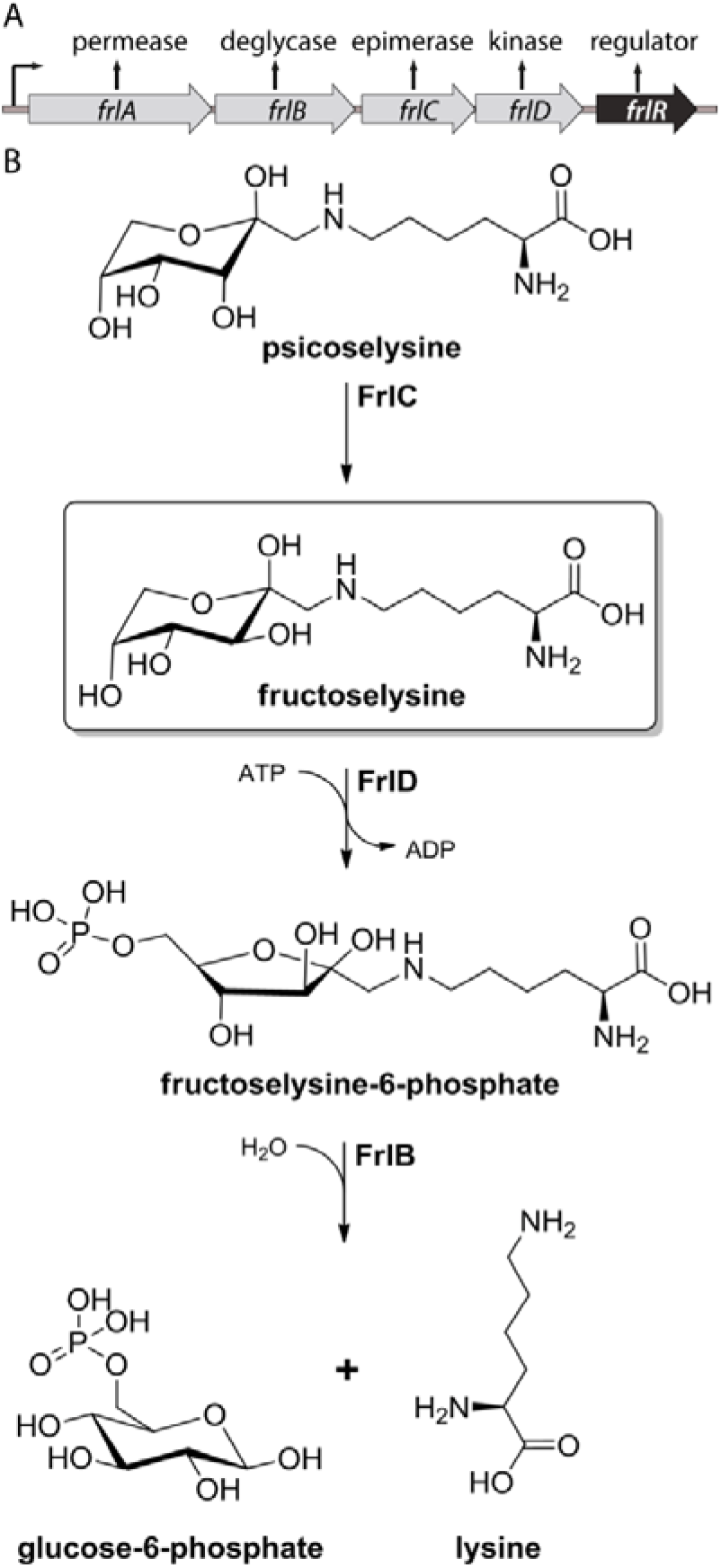
Fructose lysine metabolism in *E. coli*. A) Organisation of the *frlABCDR* genome region. B) Degradation of the ARP *N*_ε_-fructose lysine is a two-step reaction: First, a kinase FrlD phosphorylates the amino sugar, thereby forming fructose lysine-6-phosphate. In the second step, the deglycase FrlB hydrolytically cleaves sugar phosphate and amino acids into it’s the two building blocks lysine and glucose-6-phosphate, the latter of which is directly shuffled into glycolysis. A further enzyme in the pathway, termed FrlC, is capable in catalyzing the epimerization of psicose lysine into fructose lysine.

## Results

### FrlR is a putative GntR like transcriptional regulator

Wiame and coworkers noticed the presence of a putative regulator, termed FrlR, in the genomic vicinity of the ε-FrK degradation pathway (Wiame *et al.*, 2002). However, its role in controlling the *frlABCD* operon remained enigmatic. To elucidate the function of FrlR, we started with a bioinformatic comparison and performed a multiple-sequence alignment (Fig. 2). This revealed sequence similarities to the *B. subtilis* orthologous regulator FrlR of the *frlBONMD* operon (Deppe *et al.*, 2011), encoding the genes to metabolize various α-glycated amino acids (Wiame *et al.*, 2004). We also identified an ortholog in *S. enterica* - a putative regulator termed FraR - that is encoded immediately upstream of the *fraBDAE* operon, a gene cluster that is needed for degradation of fructoseasparagine (Ali *et al.*, 2014). Taken together, *E. coli* FrlR is likely to be involved in substrate-specific regulation of ARP metabolism. The outcome of our blast search also suggests a common regulatory theme that applies to all ARP metabolizing organisms despite their distinct substrate spectra.

**Fig. 2:**
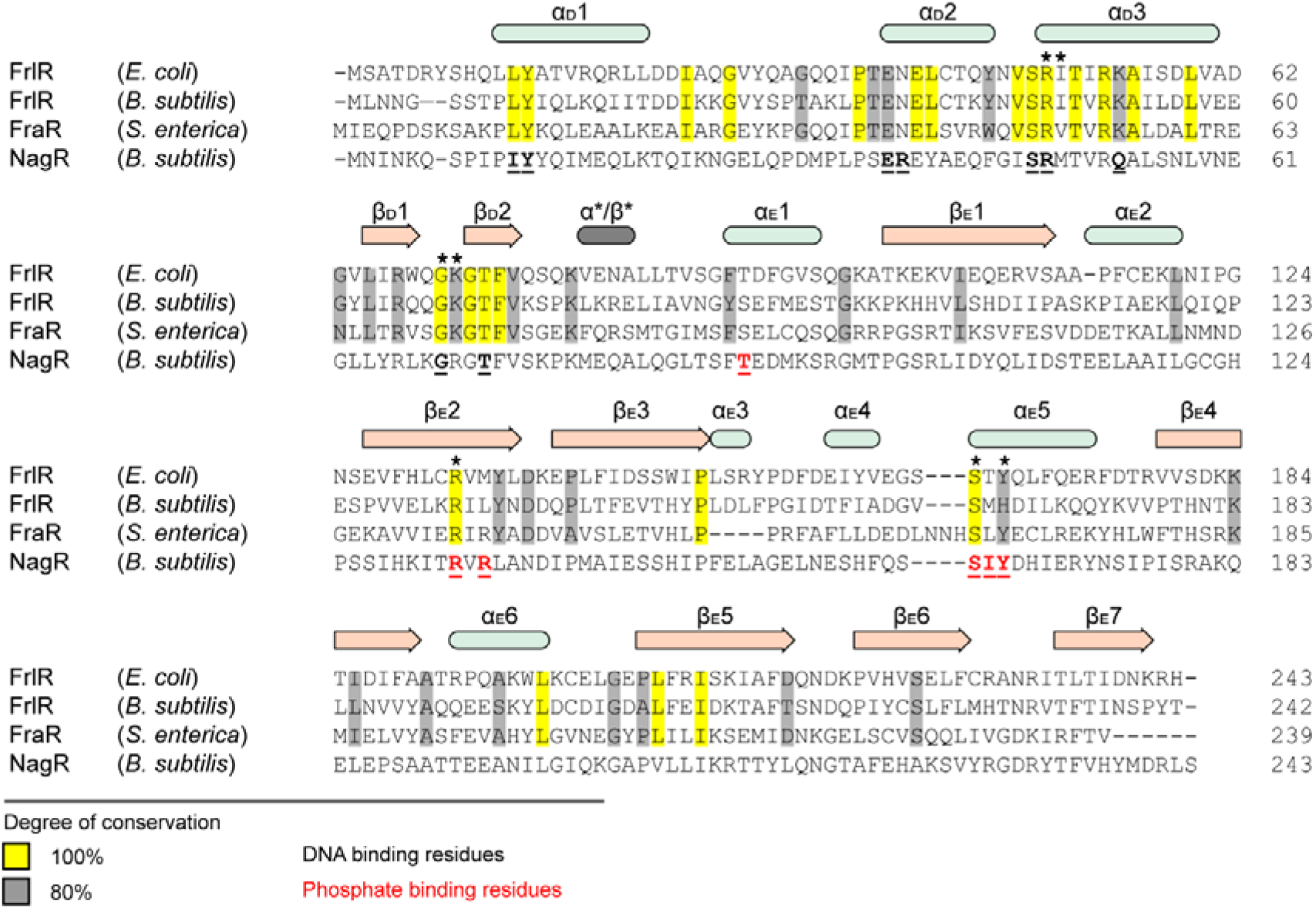
Evolutionary conservation of amino acids in FrlR homologues. Multiple-sequence alignment of FrlR/FraR proteins from *Escherichia coli, Bacillus subtilis*, *Salmonella enterica* as a selection from the alignment of 20 protein sequences that were collected from the NCBI database (Supplementary Dataset S1). The multiple-sequence alignment was generated using Clustal Omega (Sievers *et al.*, 2011). Secondary-structure elements are shown and based on the NagR crystal structure (4U0W) (Fillenberg *et al.*, 2015): α-Helices and β-strands are colored in mint and salmon, respectively. N- and C-domain elements are numbered and marked with a subscript “D” (for DNA binding domain) and “E” (for effector-binding and oligomerisation domain), respectively. Conserved residues are colored according to their degree of conservation with yellow (100 %) and grey (≥80 %). Amino acids important for function of NagR are underlined either in black - being relevant for DNA-binding - or red - being relevant for coordination of the phosphate moiety of N-acetylglucosamine-6-phosphate (Fillenberg *et al.*, 2015). “*” depict amino acids whose substitution leads to an impaired FrlR functionality.

We next used Phyre2 (Kelley *et al.*, 2015) as well as the iTAsser suite (Yang *et al.*, 2015) and performed a homology modelling to gain first molecular insights into a putative mode of action (Fig. 3). This approach revealed the cytoplasmic NagR (YvoA) of *B. subtilis* as structural homolog of FrlR (identity: 29 % / similarity: 51 %) (Resch *et al.*, 2010, Fillenberg *et al.*, 2015, Fillenberg *et al.*, 2016). NagR/DasR are GntR type transcription factors, negatively regulating the genes from the N-acetylglucosamine-degrading pathway (Resch *et al.*, 2010). Its C-terminal domain binds the effector N-acetylglucosamine-6-phosphate (GlcNAcP) and adopts a chorismate lyase fold, thus belonging to the UbiC transcription regulator-associated (UTRA) protein family (PF07702). The NagR N-terminal part, on the other hand, is a winged helix–turn–helix DNA-binding domain. It is proposed, that in an allosteric coupling mechanism GlcNAcP binding to the UTRA-domain promotes a loop-to-helix transition ultimately leading to a 122° rotation of the wHTH-domains in a so called jumping-jack-like motion (Resch *et al.*, 2010). As a result, DNA-binding is weakened and repression is abolished. Based on their structural similarities we thus speculate that NagR and FrlR might share a common molecular mechanism.

**Fig. 3:**
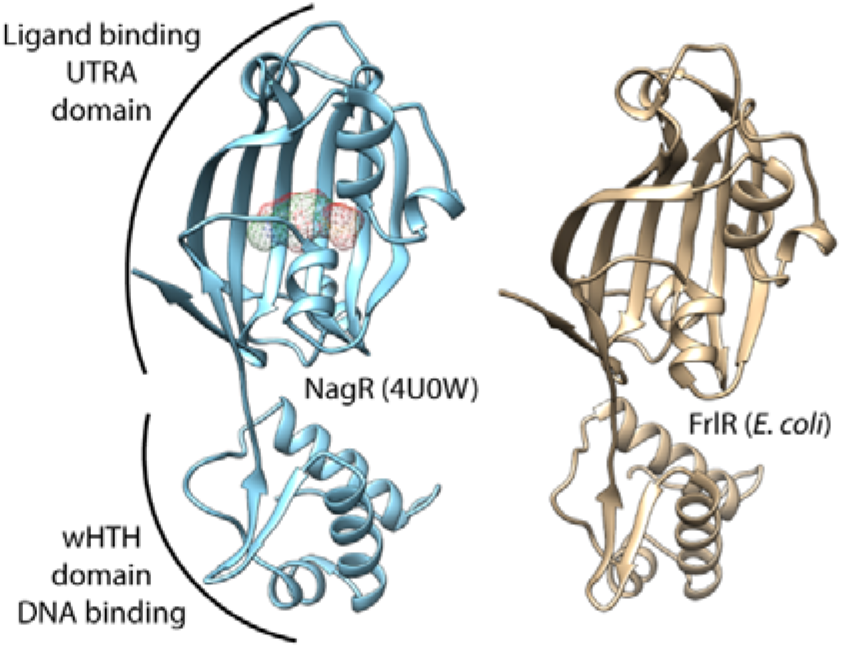
Structural organization of FrlR and comparison with NagR of *B. subtilis*. Left: Ribbon representation of the crystal structure of NagR (aa9 - aa243) of *B. subtilis* in complex with N-acetylglucosamine-6-phosphate (mesh) (4U0W). Right: Ribbon representation of the i-Tasser homology model of FrlR of *E. coli* (aa9 - aa243). Illustrations were generated with UCSF Chimera (Pettersen *et al.*, 2004).

### FrlR delays growth on fructoselysine

To investigate whether FrlR functions as a repressor similarly to NagR, we first performed a series of growth experiments with FrK. Initially, we reinvestigated the capability of *E. coli* K12 to utilize this ARP as sole carbon source. To this end, we monitored bacterial growth in minimal medium supplemented with either 1 mM of ε-FrK or 1 mM glucose, the latter serving as positive control (Fig. 4A left). As demonstrated earlier (Wiame *et al.*, 2002, Griffiths & Pridham, 1980), *E. coli* can grow on both carbon sources yielding similar biomass as can be concluded from their same maximal optical density at 600 nm. The major difference between the two curves is a prolonged lag phase and an increased doubling time of 210 min for ε-FrK supplemented cells, being around 20 % longer compared to glucose (180 min). As further control, we tested a strain Δ*frlD* lacking the FrK kinase, which catalyzes the first step in the degradation pathway. Here, growth on ε-FrK was abolished (Fig. 4A right). Reportedly, the *E. coli* FrK kinase FrlD shows only little enzymatic activity towards α-glycated amino acids *in vitro* (Wiame *et al.*, 2004) and hence we were curious whether α-FrK can substitute for the ε-glycated lysine *in vivo*. In line with the previously determined substrate specificity of FrlD, we observed neither growth with the α-glycated ARP nor any degradation (Fig. 4A left, 4B). Having confirmed these previous findings, we went on to investigate whether FrlR is involved in the regulation of ε-FrK utilization (Fig. 4B). In this regard, we compared growth of an *E. coli* wild type with a Δ*frlR* strain and found the phenotype with respect to total biomass yield indistinguishable from each other. However, at the same time we noticed a significantly shortened lag-phase, giving the first hint that FrlR acts as a repressor. In line with the assumption, the overproduction of FrlR (*frlR*^++^) prevents ε-FrK utilization. Presumably, the increased protein copy number decouples DNA binding from substrate recognition and thus reveals FrlR to be a transcriptional repressor. In parallel, we quantified metabolization of both derivatives of FrK during 24 h of incubation by direct amino acid analysis. The concentration of α-FrK remained unchanged during the experiment in the presence of the wild-type strain, whereas ε-FrK was almost completely degraded. Lysine was formed in an equimolar amount in the wild type. The results for ε-FrK were similar in the Δ*frlR* mutant. Less FrK was degraded by *frlR*^++^ cells, while no degradation and no lysine formation was observed in the Δ*frlD* strain lacking the kinase. We conclude that growth of *E. coli* is retarded when metabolization of ε-FrK is inhibited.

**Fig. 4:**
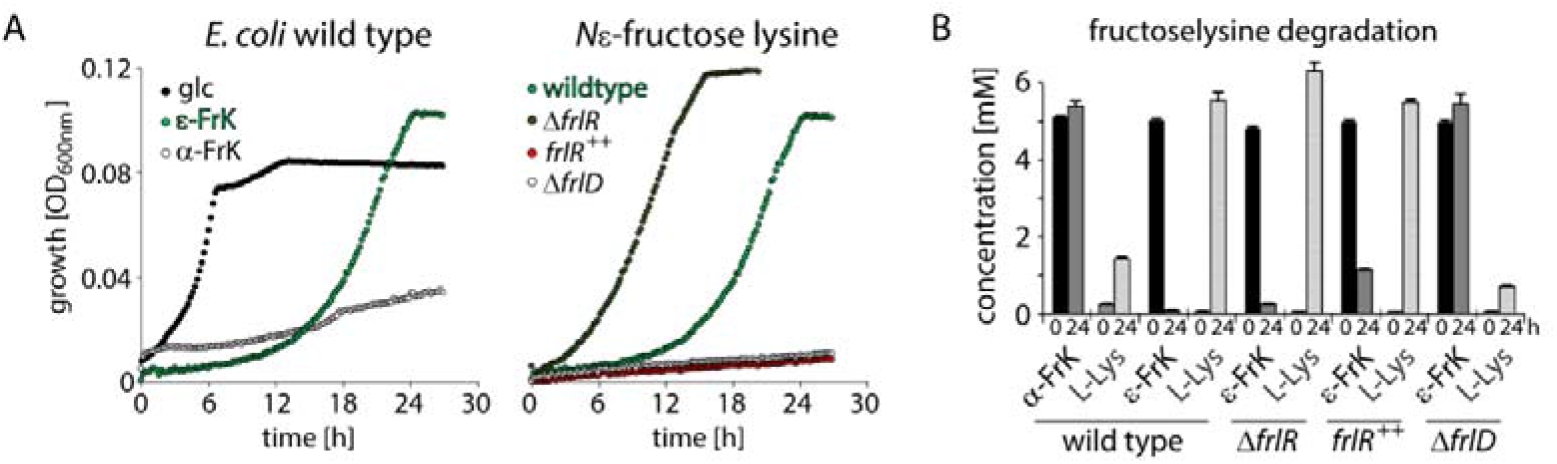
Growth analysis and fructoselysine degradation in *E. coli* wild type and *frl* mutant strains. A) Left: Growth of *E. coli* BW25113 wild type in M9-minimal medium supplemented with either 1 mM glucose (glc) (black circles), 1 mM *N*_ε_-fructoselysine (ε-FrK) (green circles) or 1 mM *N*_α_-fructoselysine (α-FrK) (grey circles). Right: Growth on 1 mM *N*_ε_-FrK as sole carbon source of *E. coli* BW25113 wild-type cells in comparison to isogenic mutant strains JW3337 (Δ*frlD*) (white circles) and JW5698 (Δ*frlR*) (black circles) (Baba *et al.*, 2006). BW25113 cells ectopically overexpressing *frlR* (*frlR*^++^) (red circles) from a pBAD33 backbone (Guzman *et al.*, 1995) was also included in the study. B) FrK degradation in *E. coli* BW25113 wild-type cells in comparison to isogenic mutant strains JW3337 (Δ*frlD*) and JW5698 (Δ*frlR*) (Baba *et al.*, 2006) as well as BW25113 cells ectopically overexpressing *frlR* (*frlR*^++^). The educts α/ε-FrK as well as the product L-lysine were quantified after 0 h and 24 h incubation time in M9-Minimal medium using cation-exchange chromatography with post-column ninhydrin derivatization and UV-detection of the reaction products (“amino acid analysis”).

### *frlABCD* transcription is positively controlled by σ32 and CRP as well as negatively regulated by FrlR

Having shown that FrK utilization is subject to FrlR regulation, we rigorously analyzed the 5’-UTR of *frlABCD* employing P_*frlABCD*_ promoter fusions to uncover further control elements. It is predicted that *frlABCD* transcription starts 75 nucleotides 5’ of the *frlA* open reading frame (+1) and depends on the housekeeping sigma factor σ^70^ (RpoD) (Fig. 5A) (Keseler *et al.*, 2017). Notably, there might be an additional initiation site 70 nucleotides further upstream recognized by the heat shock sigma factor σ^32^ (RpoH). This is especially plausible as glycation is accelerated at elevated temperatures (Hellwig & Henle, 2014). We further noticed a sequence matching the CRP/CAP (cAMP Response Protein/Catabolite Activator Protein) binding site centered around position −41.5 and thus being in perfect distance to constitute a class II promoter (Lawson *et al.*, 2004). To test our hypotheses on transcriptional regulation of the P_*frlABCD*_ promoter we initially fused 294bp (−219/+75) 5’ of *frlABCD* with the lux-operon *luxCDABE* of *Photorhabdus luminescens* (Volkwein *et al.*, 2017) and measured the light output over a time course of 24 h in *E. coli* BW25113 wild-type cells grown in Lysogeny broth (LB/Miller) (Bertani, 2004). With this, we reached a maximal luminescence per OD_600_ of about 1×10^6^ RLU showing that this region comprises a fully functional promoter (Fig. 5B-D). We note that a further sequence extension to 397bp (−322/+75) did not change this emission significantly (Fig. 5B) demonstrating that the −219/+75 lux fusion comprises all required elements for transcriptional control. The involvement of σ^32^ was investigated by ectopic overexpression of *rpoH*. This strategy was described earlier and is favored as the sigma factor is essential at growth temperatures higher than 20°C (Zhao *et al.*, 2005). With the −219/+75 P_*frlABCD*_-*lux* fusion σ^32^ overproduction led to a threefold increase in luminescence compared to the wild-type situation (Fig. 5C). At the same time lack of the putative σ^32^ dependent promoter sequence utilizing a −65/+75 fusion reduced the light output by a factor of five to 2×10^5^ RLU (Fig. 5B) both together strongly indicating for a heat stress responsive operon.

**Fig.5:**
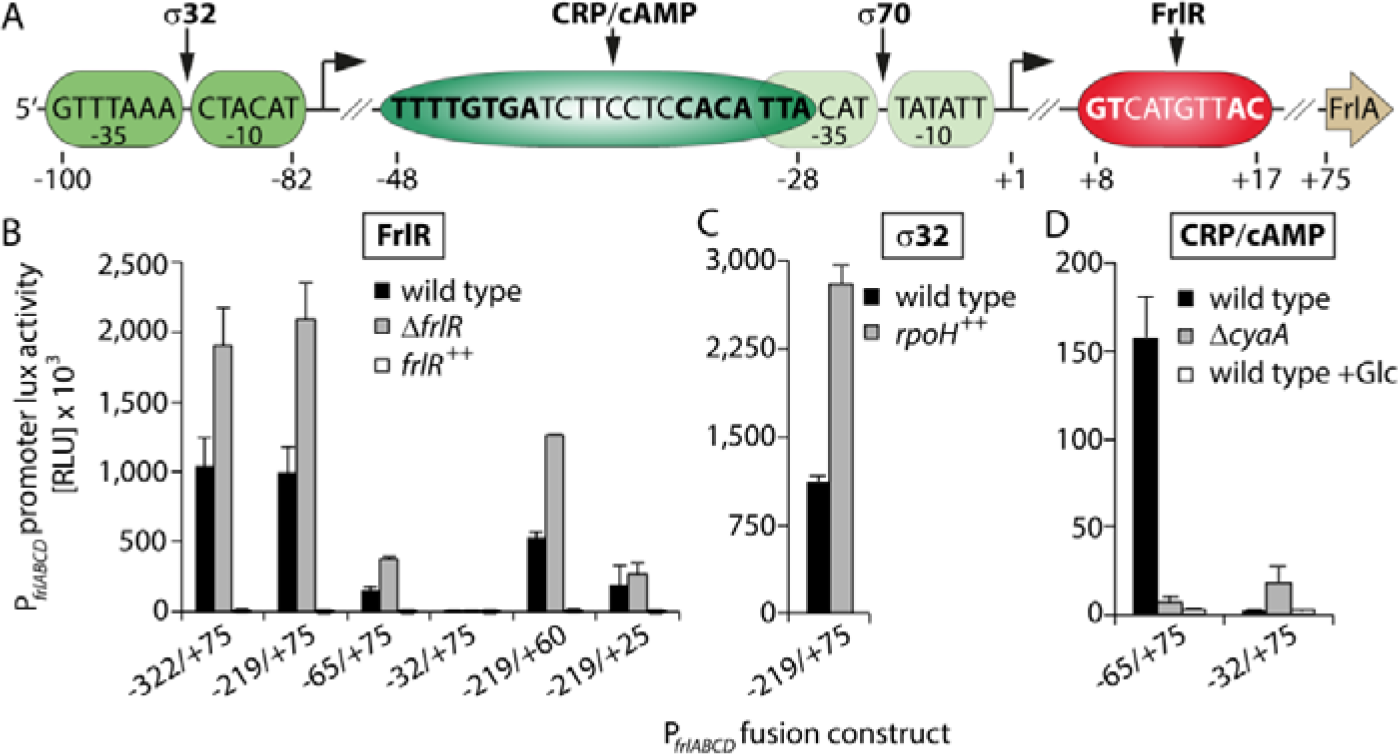
*In vivo* analysis of the *frlABCD* promoter region. A) Illustration of the *frlABCD* promoter region with +1 as the putative transcriptional start site. DNA-recognition sites for RNA-polymerase are boxed in green (dependent on RpoH/σ^32^) and light green (dependent on RpoD/σ^70^). The binding motif of the cAMP-activated global transcriptional regulator CRP is boxed in dark green. The DNA-binding site of FrlR is boxed in red. Consensus sequences for the transcription factors are highlighted with bold letters. B) - D) *In vivo* analyses of P_*frlABCD*_ promoter lux fusions in *E. coli* cells containing the respective reporter plasmid. The maximal light emission from a 24 h time course experiment of cells grown in LB (Miller) is given in RLU. The mean values of at least three biological replicates are shown. Naming of the P_*frlABCD*_-lux promoter truncations depicted on the abscissas gives information about the sequence length and numbering and relates to the illustration in A). *E. coli* BW25113 wild-type cells (black bars) in comparison to the isogenic mutant strain JW5698 (Δ*frlR*; grey bars) (Baba et al., 2006) and cells ectopically overexpressing *frlR* (*frlR*^++^; white bars) from a pBAD33 backbone (Guzman et al., 1995) were analyzed in B). C) *E. coli* BW25113 wild-type cells (black bars) in comparison to cells ectopically overexpressing *rpoH* (*rpoH*^++^; grey bars) from a pBAD33 backbone. D) *E. coli* BW25113 wild-type cells (black bars) in comparison to an isogenic mutant strain, which lacks the cAMP making adenylate cyclase CyaA, JW3778 (Δ*cyaA*). A wild-type cell culture supplemented with 20 mM glucose (glc; white bars) was also analyzed.

Next, we assessed the regulation by CRP. In an inverse correlation to the extracellular glucose concentration, this transcription factor activates metabolic processes that facilitate the metabolic conversion of alternative carbon sources. Notably, the glucose content is not measured directly but is derived indirectly from the intracellular cAMP concentration which in turn relies on the activity of a sole cAMP-synthetizing adenylate cyclase CyaA (Lawson *et al.*, 2004). Accordingly, luminescence was analyzed in the −65/+75 promoter fusion, which should comprise the full CRP DNA-binding motif. Here, LB/Miller growth conditions, with amino acids as sole carbon source, were compared on the one hand to a culture supplemented with 10mM of glucose and on the other hand to a Δc*yaA* strain (Baba *et al.*, 2006). For the latter two conditions, the light output was strongly diminished (Fig. 5D) corroborating our assumption about carbon catabolite repression. This is in line with an earlier global study that suggested CRP-dependent regulation of the *frlABCD* operon (Shimada *et al.*, 2011). We also note that glucose and cAMP dependency is lost with the truncated promoter fusion −32/+75 (Fig. 5D). However, the overall luminescence is in general strongly diminished. As this construct no longer encompasses the full CRP-binding motif, RNA-polymerase recruitment is impaired (Fig. 5A).

Lastly, we employed the Lux-reporter to investigate transcriptional regulation by FrlR. Our initial growth experiments (Fig. 4A) indicated the role of the protein as negative regulator repressing P_*frlABCD*_ under non-inducing conditions. However, the strong light emission for wild-type cells harboring P_*frlABCD*_-lux fusions in LB/Miller seems to contradict this assumption. We hypothesized, that the plasmid-borne nature of the reporter might imbalance the ratio of FrlR and promoter abundance in favor of the latter. Accordingly, increasing the FrlR copy number – by introducing an arabinose inducible plasmid-based copy of *frlR* (pBAD33-*FrlR*) – should be enough to silence the promoter and this was in fact what we observed (Fig. 5B). We also note that cells lacking *frlR* produced 1.5 fold the light of wild-type cells further supporting that FrlR is a transcriptional repressor. We ultimately analyzed whether this repression is ε/α-FrK dependent. Consequently, bioluminescence development was now measured in Δ*frlR* cells in the concomitant presence of a P_*BAD*_ controlled, plasmid-encoded copy of *frlR* (pBAD33-*FrlR*) and the −219/+75 P_*frlABCD*_-lux reporter construct. In this scenario, light emission was strongly reduced (even without arabinose induction) but becomes elevated again when adding 1 mM ε-FrK (Fig. 6). On the contrary, promoter repression was maintained with 1 mM α-FrK. These data confirm that specificity for the ε-glycated form is not limited to the metabolism, but also extends to regulation. To define the sensitivity of the system we performed a titration series with ε-FrK (10 μM-10 mM) and thus determined that the system responds to concentrations as low as 10 μM. The strength of depression is gradual with a maximum in the low mM range. Taken into account that the FrK uptake with our diet can reach such concentrations (Henle, 2005, Henle, 2003), *E. coli* is perfectly adapted to its natural human gut habitat.

**Fig. 6:**
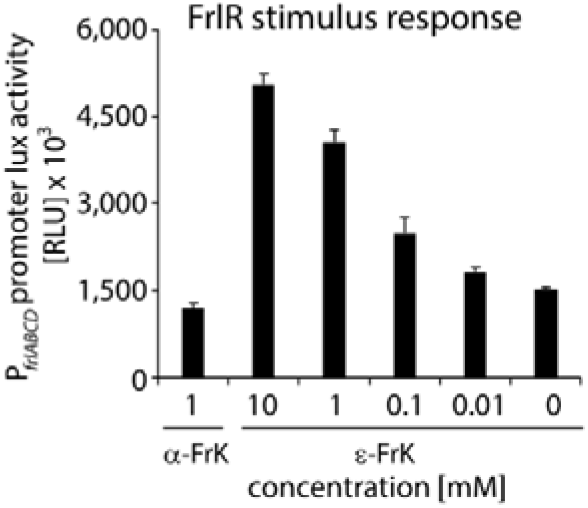
FrlR sensitivity analysis towards fructoselysine. *In vivo* analyses of the P_*frlABCD*_ promoter lux fusion −219/+75 in *E. coli* JW5698 cells (Δ*frlR*) (Baba et al., 2006) that concomitantly express *frlR* from a pBAD33 backbone (Guzman et al., 1995). The maximal light emission from a 24 h time course experiment of cells grown in LB (Miller) (without the addition of the P_*BAD*_ inducing agent L-arabinose) is given in RLU. The mean values of at least three biological replicates are shown.

### FrlR is a road block repressor that binds to a – GT(N)_6_AC - DNA consensus sequence via its wHTH-domain

Knowing that FrlR is a transcriptional repressor, we went on to identify its cognate DNA-binding motif. Therefore, we analyzed various P_*frlABCD*_-lux fusions on FrlR dependent repression (Fig. 5B). These were truncated 5’ (−322/+75, −219/+75, −65/+75 and −33/+75) or 3’ (−219/+60 and −219/+25) relative to the transcriptional start site (+1). With all tested constructs the light output decreased dramatically upon ectopic *frlR* overexpression, narrowing down the putative binding site to a region of about 50 nucleotides. The proposed structural similarities of FrlR to NagR (Fig. 3) implied that both transcription factors belong to the same family of regulators, namely the GntR/HutC related ones (Rigali *et al.*, 2002, Hoskisson & Rigali, 2009). Members of this group recognize a motif 5’-(N)_y_GT(N)_x_AC(N)_y_-3’ and so do NagR and FrlR_*Bsu*_ (Deppe *et al.*, 2011, Fillenberg *et al.*, 2015). We recognized a putative operator 5’-GT CATGTT AC-3’ immediately downstream of the transcription start site indicating that FrlR works as a road block and thus operates similar to LacI (Oehler *et al.*, 1990). To prove this hypothesis, the putative FrlR binding motif was placed between the T7-polymerase promoter (P_T7_) and a synthetic ribosome-binding site, all of which precede the ORF of the super folder green fluorescent protein (sfGFP) (Fig. 7A). An analogous construct lacking the FrlR operator (FrlO) served as control. Fluorescence intensity was measured as means of the transcriptional activity with and without FrlR. While *sfgfp* expression remained unaffected in cells with the FrlO free plasmid in all tested conditions, the presence of the DNA-binding motif strongly diminished green fluorescence intensity in an FrlR dependent manner.

**Fig. 7:**
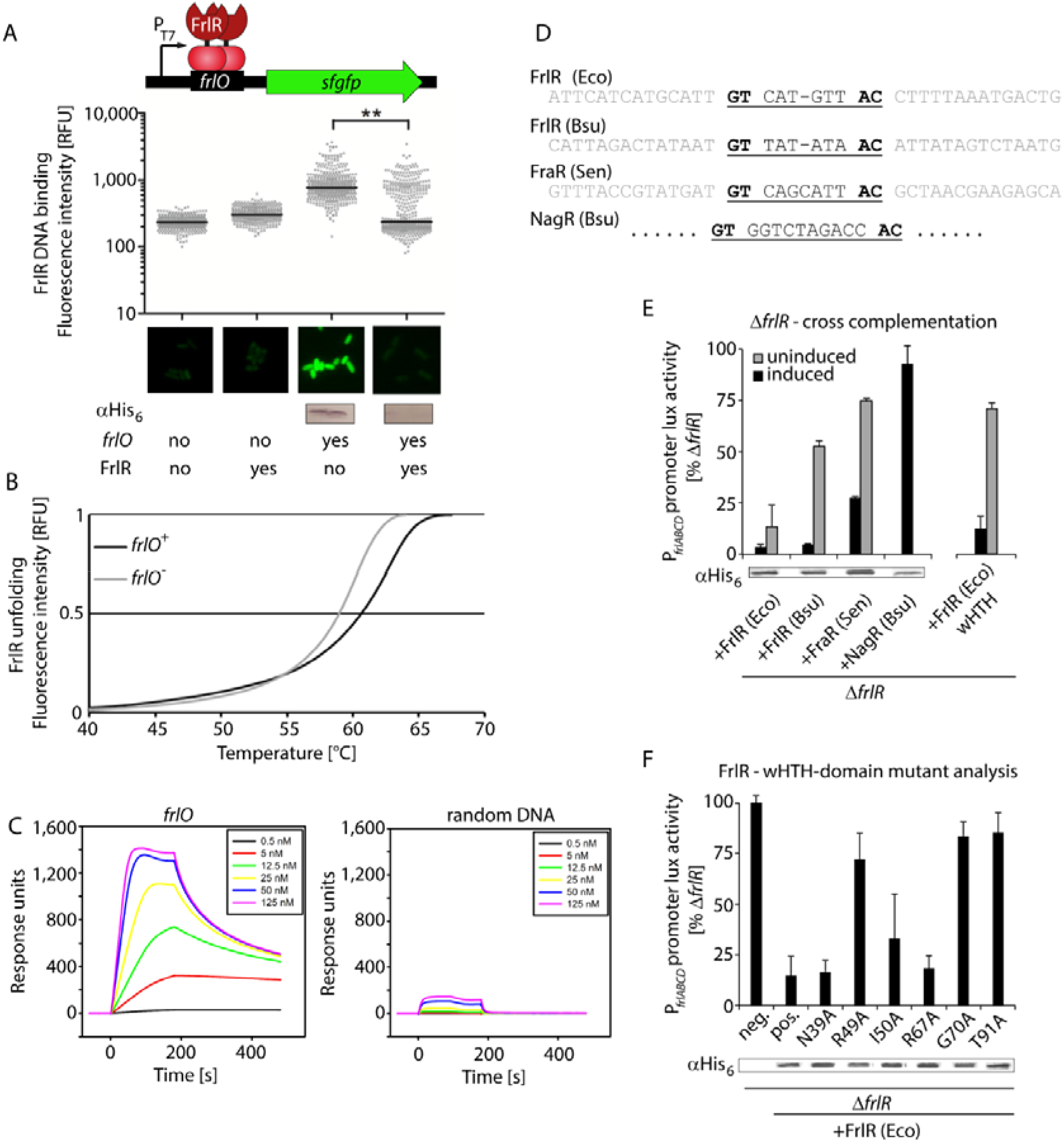
*In vivo* and *in vitro* analysis of FrlR/DNA interactions. A) FrlO dependent *gfp* expression: *E. coli* BL21 cells were cultivated in M9 minimal medium +/− arabinose and grown to the exponential growth phase. Compared are four populations all of which harbor a plasmid encoding sfGFP whose expression is driven by T7-RNA polymerase. Downstream of the T7-promoter a sequence was inserted either of a random composition (*frlO*−) or encompassing the putative FrlR recognition motif (*frlO*+). Depicted is the fluorescence intensity (given in relative fluorescence units RFU) of 300 individual cells in the presence or absence of FrlR (FrlR+/−). Pictures were analyzed using ImageJ (Schneider *et al.*, 2012). sfGFP-production was visualized by Western blot analysis using anti-His_6_ specific antibodies (α-His_6_). B) *In vitro* DNA binding of FrlR to *frlO* analyzed by thermal shift assays (TSA): TSAs were conducted for FrlR in the presence of a DNA fragment either comprising *frlO* (black line) or a random sequence (grey line). Depicted are the FrlR melting curves as a function of fluorescence with 0 (fully folded) and 100 (fully denatured). Temperature was increased 0.5 °C every 30 seconds from 8°C to 90 °C and fluorescence was monitored in each step. C) *In vitro* DNA binding of FrlR to P_*frlO*_ analyzed by surface plasmon resonance spectroscopy (SPR): The biotin-labeled DNA fragment *frlO* (left), and the control fragment without the FlrR binding motif (right) were captured on a streptavidin-coated sensor chip, and purified FlrR was passed over the chip at a flow rate of 30 μl/min and temperature of 25°C [concentrations of 0.5 nM, 5 nM, 12.5 nM, 25 nM, 50 nM, 125 nM), using a contact (association) time of 180 sec, followed by a 300-sec dissociation phase. The increase in RU correlates with an increasing FlrR concentration. D) Comparison of the (putative) FrlR, (FraR), NagR DNA-binding motifs from *E. coli* (Eco), *B. subtilis* (Bsu) and *S. enterica* (Sen). E) Left: repressor activity on the *E. coli* FrlO operator by FrlR or from *E. coli* (Eco) and *B. subtilis* (Bsu) as well as FraR of *S. enterica* (Sen). Right: repressor activity by the *E. coli* FrlR winged helix-turn-helix DNA-binding domain (wHTH). Experiments were conducted in *E. coli* JW5698 cells (Δ*frlR*) (Baba et al., 2006) encoding a P_*BAD*_-inducible copy of the respective gene or gene fragment. Depicted is the maximal light emission from a 24 h time course experiment of cells grown in LB (Miller) with (black bars) and without (grey bars) the addition 0.2 % L-arabinose. The mean values of at least three biological replicates are shown. Protein production was confirmed by Western blot analysis using anti-His_6_ specific antibodies (α-His_6_) F) FrlR-wHTH-domain mutant analysis. Assessed is the capability of FrlR mutant variants to repress the *frlABCD* promoter lux activity. Experiments were conducted in *E. coli* JW5698 cells (Δ*frlR*) (Baba et al., 2006) encoding a P_*BAD*_-inducible copy of the respective mutants. Depicted is the maximal light emission from a 24 h time course experiment of cells grown in LB (Miller). Neg.: Δ*frlR* Pos.: Δ*frlR*+FrlR^wt^. Protein production of the FrlR variants was confirmed by Western blot analysis using anti-His_6_ specific antibodies (α-His_6_).

Our *in vivo* analysis was complemented by testing the operator sequence on FrlR binding *in vitro.* We employed thermal shift assays (Huynh & Partch, 2015) to assess heat stability of FrlR in the presence of *frlO* (*frlO^+^*) or a random DNA-fragment (*frlO^−^*) of similar size (Fig. 7B). In theory, DNA-binding to FrlR should stabilize the protein and in turn, the melting temperature increases. In fact, we observed a FrlR melting temperature of 62°C in combination with *frlO*, whereas it was 3°C lower in the mixture with the random DNA-fragment.

As these data are only qualitative, we next employed Surface Plasmon Resonance spectroscopy (SPR) in order to determine the binding kinetics (association rate *k*_a_ and dissociation rate *k*_d_) as well as the affinity (K_D_) of FrlR and its cognate recognition motif (Fig. 7C, Fig. S1). To this end, different concentrations of FrlR were combined with two immobilized DNA fragments (see the experimental procedures for details). One fragment includes the FrlR binding site, the other was of random sequence composition and served as negative control. Before that, the absolute and “active” fraction of FrlR was determined using calibration free concentration analysis (CFCA) to approximately 50 % of the total protein concentration. A clear and stable binding could be observed for FrlR to *frlO* with a high association (*k*_a_=7.0 × 10^6^ M^−1^s^−1^) and low dissociation (*k*_d_=3.4 × 10^−2^ s^−1^) rate, whereas only a weak binding of FlrR was seen with the control DNA. Calculations were based on the association and dissociation rates and with this, we derived a high affinity of FrlR for *frlO* of 4.9 nM.

We were also curious whether the *E. coli* FrlR binding motif is recognized by the orthologous regulators FrlR of *B. subtilis* and FraR of *S. enterica.* In this regard, the corresponding genes were placed under control of the arabinose inducible promoter P_*BAD*_ analogous to *frlR* of *E. coli*. NagR of *B. subtilis* was also included in the study and served as negative control. Binding to 5’-GT CATGTT AC-3’ was tested once more employing the P_*frlABCD*_-lux reporter. Strikingly, despite the significant differences in sequence composition of their cognate DNA-binding motifs (Fig. 7D), the two proteins FrlR_*Bsu*_ and FraR but not NagR negatively regulated P_*frlABCD*_ dependent *luxCDABE* expression. We hypothesized therefore that rather the spacer length between the flanking GT and AC – which are six base pairs (bp) in case of FrlO_*Eco*_ - is most important. This would explain not only the lack of repression with NagR (10 bp spacer length) but also the more subtle differences we observed when comparing the light output of cells encoding FrlR with those harboring *frlR_Bsu_* and *fraR*. Whereas a leaky *frlR* expression (no arabinose for promoter activation added) is sufficient for efficient silencing, FraR with its presumed seven bp long spacer completely lost its regulatory capability under non-inducing conditions. On the contrary also FrlR_*Bsu*_ did repress FrlO_*Eco*_, which is plausible as FrlO_*Bsu*_ also encompasses six bp. The reduced repression efficiency compared to *E. coli* FrlR can be explained by the differences in sequence composition. In conclusion, our data supports a model in which the length of the GT/AC bracketed spacer predominately defines the common theme for operator recognition while the motif composition modulates its strength. To assess DNA binding by FrlR, we first verified DNA binding by the putative wHTH-DNA binding domain (Fig. 3). DNA-binding to the recognition sequence can be enforced by overproduction of the wHTH-domain solely (Schlundt *et al.*, 2017). Accordingly, we truncated FrlR and compared the fragment encompassing amino acids 1-77 with the full-length protein (244 aa). In *E. coli* cells, that simultaneously harbor the P_*frlABCD*_-*lux* −219/+75 reporter plasmid luminescence can be suppressed only when transcription of the FrlR 1-77 aa fragment was arabinose induced while with the full-length FrlR even low protein levels diminished the light output (Fig. 7E). These data strongly imply that indeed the first 77 residues fold into a wHTH-domain that is sufficient to recognize the promoter. However, full-length FrlR is needed for high affinity binding.

Next, we generated FrlR mutants based on the sequence alignment with NagR (Fig 2). In NagR, the wing motif reaches into the minor groove and contacts the flanking guanosine via the carbonyl oxygen of Gly69 (Fillenberg *et al.*, 2015). In addition, the two conserved arginines Arg38 and Arg48 specifically recognize further guanosines of the operator, by forming bidental contacts with their corresponding guanine base. Taken together, these three amino acids are assumed to be the hallmark feature of the specific interaction of NagR with DNA (Fillenberg *et al.*, 2015). Two of the contacts - Gly69 and Arg48 - are also conserved throughout FrlR/FraR homologs (Fig. 2). Accordingly, we mutated the corresponding residues – Arg49 and Gly70 – into alanine resulting in the FrlR variants R49A and G70A. We additionally mutated the putative nucleic acid interacting residues N39, I50, R67 and K71 into alanine, being in the equivalent positions to NagR’s Arg38, Met49 and Arg70, respectively. The functionality of the FrlR variants was tested *in vivo* on their capability to abolish luminescence in Δ*frlR* cells encoding the −219/+75 P_*frlABCD*_-lux reporter. Three out of the six mutants – R49A, G70A and K71A – clearly lost their repressing capability, indicating their role in DNA binding. This led us to conclude that FrlR binds to its recognition motif in a way similar as described for NagR.

### FrlR dimerizes via its C-terminal UTRA-domain

The predicted structural similarities to NagR combined with our data on FrlR DNA-binding implies that substrate binding might also occur analogously. Canonically, members of the GntR/HutC family of transcriptional regulators form antiparallel dimers via their UTRA domain to accommodate the two half sites of their palindromic GT/AC flanked DNA-binding motif (Rigali *et al.*, 2002, Fillenberg *et al.*, 2015, Suvorova *et al.*, 2015). Whether this mode of action also applies for FrlR we examined the dimerization tendency *in vitro* and *in vivo*. For the *in vivo* analysis, the bacterial-two-hybrid system described by Karimova et al. was employed (Karimova *et al.*, 1998) with the exception that all experiments were carried out with our recently published reporter strain, that has a luminescence reporter readout in addition to the original LacZ based colorimetric one (Volkwein *et al.*, 2019). Based on the FrlR homology modeling (Fig. 3) we split the protein into two parts, one comprising the N-terminal DNA-binding domain (aa 1-77), the other one encompassing the C-terminal UTRA-domain (aa 78 - aa 244) and fused them to the T25 and T18 fragments of *Bordetella pertussis* adenylate cyclase. Bioluminescence was recorded qualitatively on agar plates together with the GCN4 leucine zipper as positive (bright phenotype) and T25/T18 solely as negative control (dark phenotype) (Fig. 8A). When assessing the self-interaction of the UTRA-domain we saw the bright phenotype, whereas clones with the wHTH-domain remained dark, confirming our initial assumption. We also tested whether FrK influences interaction strength or might even interfere with dimer formation. Consequently, we measured bioluminescence development quantitatively by recording the light output in a 24 h time course experiment. The maximal RLU are plotted as a bar diagram (Fig. 8B). Neither the addition of the α-nor the ε-glycated FrK abolished light emission. However, we observed a slight but significant change in interaction strength with the latter. Reportedly, NagR derepression is not achieved by monomerization upon signal perception (Resch *et al.*, 2010) as e.g. in the case of other one-component systems such CadC (Lindner & White, 2014, Buchner *et al.*, 2015). Instead, substrate binding to the UTRA-domain is transduced into a structural rearrangement of the two wHTH-domains, that hinders proper DNA-binding. Similarly, such movement might also reposition the two fragments T18 and T25 and could explain the reduction in bioluminescence with ε-FrK. This is in so far plausible as the loop, which is transitioned into a helix upon N-acetylglucosamine binding to NagR (Resch *et al.*, 2010, Fillenberg *et al.*, 2015) was kept in our fusion construct.

**Fig. 8:**
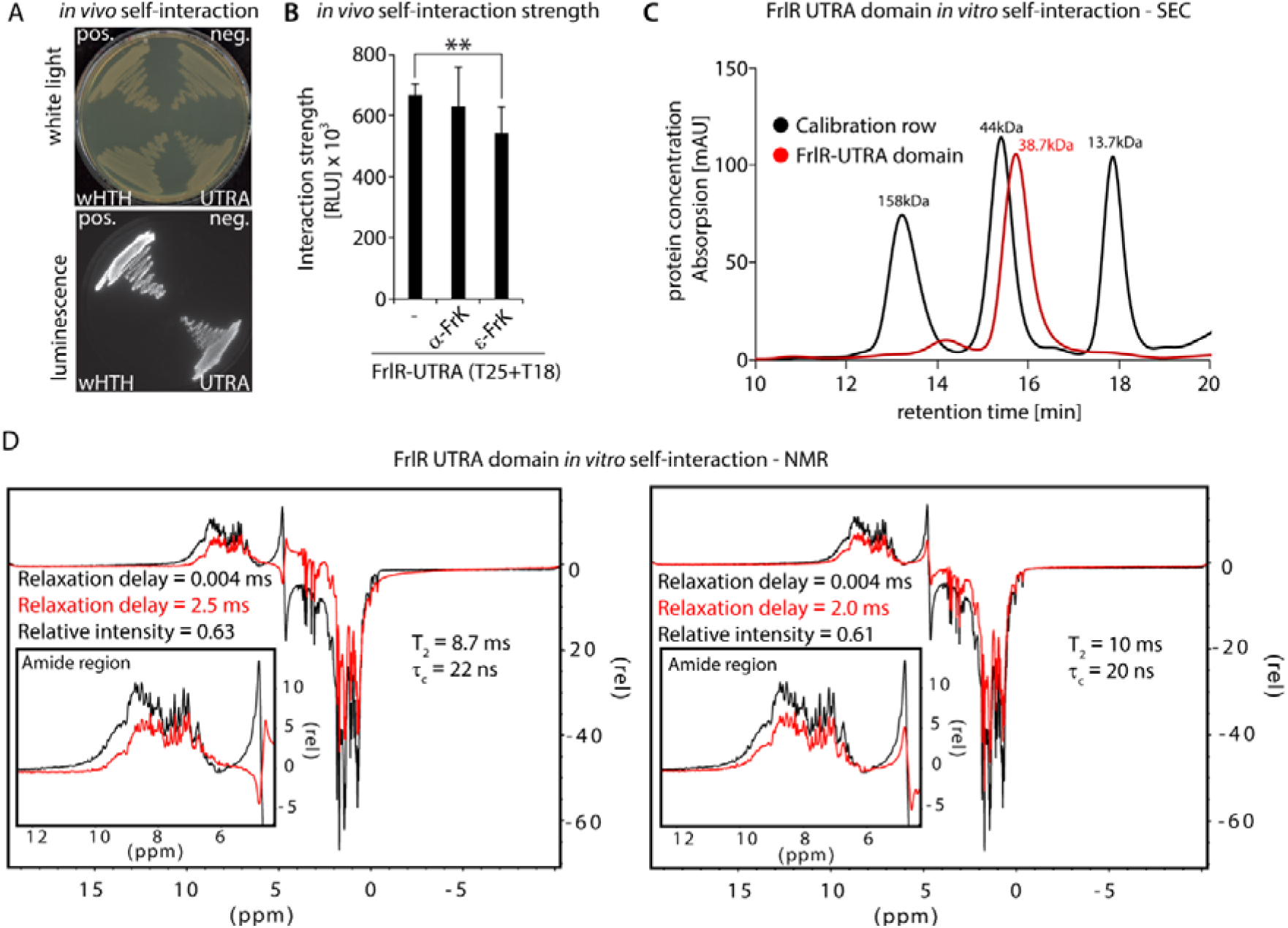
FrlR domains *in vivo* / *in vitro* interaction analyses. A) Qualitative *in vivo* self-interaction analysis of a T18/T25-FrlR UTRA-domain fusion as well as a T18/T25-wHTH-domain fusion. B) Quantitative *in vivo* self-interaction analysis of a T18/T25-FrlR UTRA-domain fusion with and without (-) supplement of α-FrK or ε-FrK. The maximal light emission from a 40 h time course experiment is given in RLU. 95 % confidence intervals of at least six replicates are shown. Asterisks indicate significant (P < 0.05) differences in the maximal light emission between cells without supplement and those being exposed to ε-FrK. C) Size exclusion chromatography of purified FrlR UTRA-domain. The elution profile is depicted as solid red line. The black line represents a calibration curve of three proteins with varying size (158 kDa, 44 kDa and 13.7 kDa). Based on this the FrlR UTRA-domain migration behavior corresponds to a size of 38.7 kDA. D) 1D-jump-and-return NMR experiments including 0.004, 2.0 and 2.5 ms relaxation delays from which the transverse relaxation time *T*_2_ is estimated to be 9 ms, allowing to infer a rotational correlation time τ_c_ of 21 ns assuming isotropic tumbling. This indicates FrlR to be a stable dimer in solution.

To recapitulate our *in vivo* observation *in vitro*, we investigated the oligomerization behavior of the UTRA domain by determining its molecular weight by size exclusion chromatography (Fig 8C). In its monomeric form, this would correspond to 19.3 kDa. As expected and in line with the BTH data, FrlR_UTRA_ runs at the size of dimers. The addition of ε-FrK did not change the migration behavior of FrlR_UTRA_ (data not shown) further supporting that the regulation does not alter the oligomeric state of the protein. In addition to this, nuclear magnetic resonance (NMR) spectroscopy also confirmed the dimeric state of the FrlR_UTRA_ domain. The line width of ^1^H-^15^N resonances in ^1^H-^15^N-HSQC spectra is larger than what would be expected for a monomeric FrlR_UTRA_ (Fig. S2A). Receptor-based titration with ε-FrK did not change the resonance line width in ^1^H-^15^N-HSQC spectra and we could only observe small chemical shift perturbations (Fig. S2A) even with 6-fold excess of ε-FrK, indicating that interaction with ε-FrK is weak and FrlR_UTRA_ does not become monomeric upon titration with ε-FrK. This qualitative assessment of FrlR_UTRA_ dimerization was followed up with 1D *T_2_* experiments, in which a relaxation delay has been included in a 1D-^1^H experiment to estimate the transverse relaxation time *T_2_*. From three different relaxation delays (0.004, 2.0 and 2.5 ms) we could estimate *T_2_* to be 9 ms (Fig. 8D). Assuming isotropic tumbling, we estimate a rotational correlation time of 21 ns which would correspond to a molecular weight of 42 kDa. This indicates that FrlR tumbles as a dimer in solution. The weak interaction with ε-FrK is confirmed by ligand-based titration using saturation transfer difference (STD)-NMR, where no magnetization transfer could be observed from protein to ligand suggesting an affinity weaker than 10^−3^ M for the FrlR-ε-FrK interaction (Fig. S2 B, C).

### Fructoselysine-6-phosphate is the cognate effector substrate of FrlR

We have conclusively shown, that ε-FrK efficiently induces *E. coli* FrlR dependent derepression *in vivo*. However, our *in vitro* FrlR interaction analysis with the ligand showed only a weak response (Fig. 6, SX) or even failed (Fig. S2B, C). Thus, we hypothesized, that not FrK directly but instead one of its metabolic derivatives is the cognate substrate. This might be a plausible explanation as e.g. the structural homolog of FrlR – NagR – recognizes the phosphorylated form of GlcNAc, *N*-acetylglucosamine-6-phosphate (Resch *et al.*, 2010, Rigali *et al.*, 2002). Analogous to this, the most probable candidate is ε-FrK-6-phosphate, which is generated by the kinase FrlD as first step of the degradation pathway (Fig. 1). If the assumption is true, loss of FrlD will abolish ε-FrK mediated derepression of P_*frlABCD*_. We therefore utilized an *E. coli* Δ*frlD* strain containing the −219/+75 P_*frlABCD*_-*lux* fusion and measured the light output in the presence and the absence of the ARP. Indeed, the *frlD*^−^ strain lost its ability to respond to ε-FrK, as can be inferred from the maximum light emission, which – unlike *frlD*^+^ cells – did not increase with external supplementation of the ARP (Fig. 9).

**Fig. 9:**
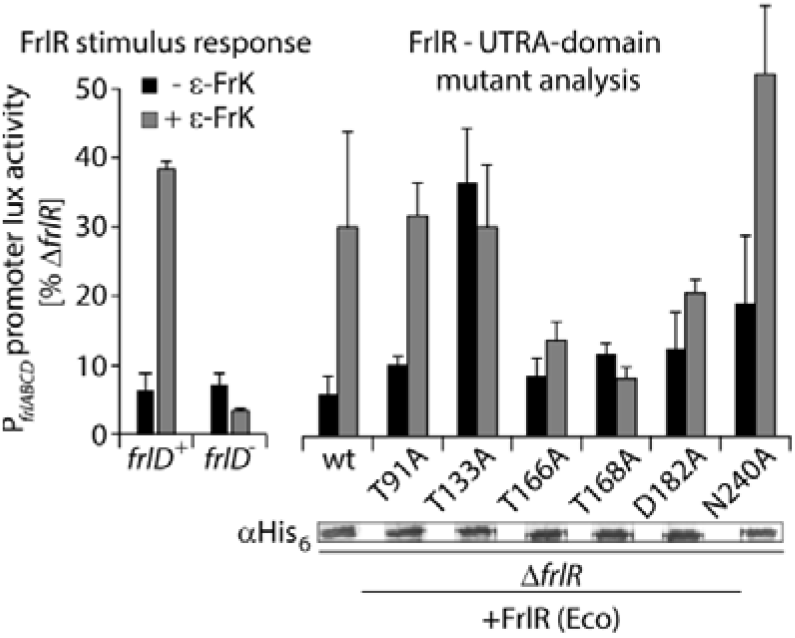
FrlR mutant analysis towards FrK-6P. *In vivo* analyses of the P_*frlABCD*_ promoter lux fusion − 219/+75 in *E. coli* JW5698 cells (*frlD*^+^) and JW3337 (*frlD*^−^) (Baba et al., 2006) that concomitantly express *frlR* and mutant derivatives from a pBAD33 backbone (Guzman et al., 1995). The maximal light emission from a 24 h time course experiment of cells grown in LB (Miller) (without the addition of the P_*BAD*_ inducing agent L-arabinose) is given in RLU. The mean values of at least three biological replicates are shown. Protein production of the FrlR variants was confirmed by Western blot analysis using anti-His_6_ specific antibodies (α-His_6_)

The sequence comparison of FrlR and NagR revealed high conservation of the phosphate-binding pocket (Fig. 2). Therefore, we decided to investigate, whether ε-FrK-6-phosphate might be accommodated analogously as N-acetylglucosamine-6phosphate in NagR (Fig. 3). Accordingly, we generated the substitution variants located in the UTRA domain, namely T91A, T133A, T166A, T168A, D182A and N240A, and investigated them on their capability to regulate promoter activity in an ε-FrK dependent manner once more utilizing the −219/+75 P_*frlABCD*_-lux fusion. With the wild-type protein and in the absence of ε-FrK we observed a strong repression, exhibited by an about 95 % diminished light output compared to cells lacking *frlR* (Fig. 9). The luminescence increased by more than 5-fold when the culture was supplemented with the ARP. Deviating from this behavior T166A, T168A, and D182A did not respond to the presence of ε-FrK with P_*frlABCD*_ remaining repressed. Similarly, T33A became blind to the inductor but at the same time also lost its DNA-binding capability. We hypothesize, that this variant is locked in its inactive state that precludes efficient operator recognition. In summary, these data show that ε-FrK-6-phosphate is the cognate substrate of FrlR and recognizes similarly as NagR.

## Discussion

In the present study, the transcriptional regulation of FrK metabolism in *E. coli* was investigated. Our data show that the expression of the *frlABCD* operon is controlled by global and specific stimuli. We identified two promoter regions, one driven by the alternative sigma factor RpoH (σ32). The induction of the other σ70-dependent promoter is subject to cAMP/CRP triggered catabolite repression. A road block repressor FrlR further prevents transcription in the absence of FrK. However, this substrate has to be processed to fructoselysine-6-phosphate by the kinase FrlD to be recognized as the cognate stimulus. Accordingly, a basal *frlABCD* gene expression is necessary even under repressive conditions. Several scenarios are conceivable to achieve such a goal. For instance, the copy number of the FrlR protein could be limited to such an extent that not every cell has a sufficient amount for effective repression. In this case, an unequal distribution of the permease FrlA and kinase FrlD occurs within the population, which enables a heterogeneous response to the carbon source. The resulting heterogeneity might be particularly pronounced because of the complex regulation and the necessity to transport and modify FrK to induce the cell response. The so-called “bet hedging” strategy (van Vliet and Ackermann, 2015) would be particularly advantageous here: Limiting the FrK metabolism to a subset of cells allows the simultaneous utilization of different carbon sources by the entire population. In the intestine, *E. coli* is confronted with exactly such a situation and is thus able to compete with other bacteria of the gut microbiota. In the same context, the specialization of the metabolism towards certain MRPs is also beneficial. While *E. coli* exclusively utilizes ε-fructose/psicoselysine, *B. subtilis* favors various α-glycated amino acids (Wiame *et al.*, 2004, Wiame & Van Schaftingen, 2004), thus creating distinct niches and minimizing competition. However, such selectivity also demands for species specific regulation patterns. Global transcriptional control seems to be adjusted to the respective organism with *B. subtilis* being dependent on CodY, a transcriptional regulator that helps to adapt to changes in nutrient availability (Deppe *et al.*, 2011) and *E. coli* where MRP metabolism is activated by CRP/cAMP and σ^32^ instead. On the contrary, the substrate specific FrlR proteins from both organisms are 63 % similar and even recognize the same operator sequence. It is therefore possible that both FrlRs also respond to the same stimulus; however, in our assays we saw a highly specific response of *E. coli* FrlR towards ε-FrK. ε-FrK-6P is the cognate signal and thus an elegant way to achieve regulatory specificity is metabolic specificity. The FrlD kinases from *E. coli* and *B. subtilis* differ in their enzymatic properties (Wiame *et al.*, 2004). The K_M_ of ε-FrK for instance is 20 μM with FrlD_*Eco*_, the corresponding one for FrlD_*Bsu*_ (YurL) is three orders of magnitude lower (14 mM). Conversely, α-fructosevaline is processed efficiently by the *B. subtilis* enzyme at a concentration of about 100 μM, whereas the *E. coli* counterpart has a K_M_ above 20 mM. The selectivity of the metabolism can also explain how FrlR is able to respond to chemically diverse α-glycated substrates. Specifically, substrate recognition could simply be achieved by the combination of sugar 6-phosphate and C_1_-N linkage, two structure elements shared by all phosphorylated ARPs derived from glucose.

## Experimental Procedures

### Bacterial strains, plasmids and growth conditions

Bacterial strains and plasmids and primers are listed in Tables S1-3. *E. coli* was routinely cultivated in LB according to the Miller modification (Bertani, 1951, Miller, 1992) or M9 minimal medium (Miller, 1972) unless indicated otherwise. For solidification 1.5 % (w/v) agar was added to the medium. If needed, carbon sources and other media supplements were added as indicated. Antibiotics were used at the following final concentrations: 100 μg/ml ampicillin sodium salt, 50 μg/ml kanamycin sulfate, 30 μg/ml chloramphenicol, or 20 μg/ml gentamycin sulfate). Growth was recorded by measuring the optical density at a wavelength of 600 nm. Plasmids carrying the pBAD (Guzman *et al.*, 1995) or T7 promoter were induced with L-arabinose at a final concentration of 0.2 % (w/v) or Isopropyl-β-D-thiogalactopyranosid (IPTG) at a final concentration of 1mM.

All kits and enzymes employed for plasmid construction were used according to manufacturer’s instructions: Plasmid DNA was isolated using the Hi Yield® Plasmid Mini Kit from Süd-Laborbedarf GmbH. DNA fragments were purified from agarose gels using the Hi Yield® Gel/PCR DNA fragment extraction kit from Süd-Laborbedarf GmbH. All restriction enzymes, DNA modifying enzymes and the Q5® high fidelity DNA polymerase for PCR amplification were purchased from New England BioLabs GmbH. A detailed description for plasmid construction is given in Table S2.

### Synthesis and analysis of ARPs

N-ε- as well as N-α-FrK were synthesized and isolated according to previous publications (Krause *et al.*, 2003, Hellwig *et al.*, 2011) and met the spectroscopic properties given in those works. Analysis of both ARPs as well as lysine was performed by amino acid analysis with the analyzer S 433 (Sykam) on a cation-exchange column (LCA K07/Li; 150 mm × 4.6 mm, 7 μm). Before analysis, 10 μL of solutions from microorganism culture was diluted with 190 μL of loading buffer (0.12 M lithium citrate, pH 2.12), centrifuged (10.000 x g, 10 min), and 40 μL of the diluted sample was injected. Separation was accomplished with custom lithium citrate buffers of increasing pH and ionic strength. Amino acids were detected by online post-column derivatization with ninhydrin and UV-detection (570 nm). Calibration was performed by an external standard of proteinogenic amino acids, and both FrK derivatives were quantified as lysine (Hellwig *et al.*, 2011).

### SDS-PAGE and Western Blotting

For protein analyses cells were subjected to 12.5 % (w/v) sodium dodecyl sulfate (SDS) polyacrylamide gel electrophoresis (PAGE) as described by Laemmli (Laemmli, 1970). To visualize proteins by UV light 2,2,2-trichloroethanol was added to the polyacrylamide gels (Ladner *et al.*, 2004). Subsequently, the proteins were transferred onto nitrocellulose membranes, which were then subjected to immunoblotting. In a first step the membranes were incubated either with 0.1 μg/mL Anti-6×His^®^ antibody (Abcam) to detect EF-P, or with 0.1 μg/ml Anti-GFP (Abcam) antibody. These primary antibodies (rabbit) were targeted with 0.2 μg/ml Anti-rabbit alkaline phosphatase-conjugated secondary antibody (Rockland) and detected by adding development solution [50 mM sodium carbonate buffer, pH 9.5, 0.01 % (w/v) p-nitro blue tetrazolium chloride (NBT) and 0.045 % (w/v) 5-bromo-4-chloro-3-indolyl-phosphate (BCIP)].

### Bacterial Two-Hybrid Assay

Protein-protein interactions were detected using the bacteria adenylate cyclase two-hybrid system kit (Euromedex) according to product manuals (Karimova *et al.*, 2000). This system is based on functional reconstitution of split *Bordetella pertussis* adenylate cyclase CyaA, which catalyzes the formation of cyclic AMP from ATP. In *E. coli* KV1, the cAMP depedent *lac* promoter P_*lac*_ precedes a translational fusion of the *lux*-operon and *lacZ*, allowing the indirect measurement of protein-protein-interactions by light emission and colorimetric detection (Volkwein *et al.*, 2019).

For measuring interaction strength, chemically competent (Inoue *et al.*, 1990) *E. coli* KV1 cells were transformed with pKT25-*frlR* FL, pKT25-*frlR* UTRA-domain or pKT25-*frlR* HTH-domain and pUT18C-*frlR* FL, pUT18C-*frlR* UTRA-domain or pUT18C-*frlR* HTH-domain. Transformants containing pUT18-*zip*/pKT25-*zip* and pUT18C/pKT25 vector backbones were used as positive and negative controls, respectively. Single colonies were inoculated in LB (with 50 μg/ml kanamycin sulfate and 100 μg/ml ampicillin sodium salt, 5 mM α-/ε-FrK and 0.5 mM IPTG (w/v)) and grown aerobically at 37 ° o/n. The next day, a microtiter plate with fresh LB (with the appropriate antibiotics, 10 mM substrate and 0.5 mM IPTG (w/v)) was inoculated with the cells at an OD_600_ of 0.01. The cells were grown aerobically in the Tecan Infinite F500 system (TECAN) at 30°C. OD_600_ and luminescence were recorded in 10 min intervals over the course of 16 h. Each measurement was performed at least in triplicate.

For qualitative analysis, the bacterial two-hybrid KV1 strains were plated on LB agar (containing 50 μg/ml kanamycin sulfate, 100 μg/ml ampicillin sodium salt and 0.5 mM IPTG) and grown overnight at 37 °C. Pictures of the plates were taken in a Fusion-SL 3500 WL (PEQLAB) using 10 seconds of exposure time.

### Luminescence activity assay

Single colonies were inoculated in LB (with appropriate antibiotics and 0.2 % arabinose (w/v)) and grown aerobically at 37 °C. The next day, a microtiter plate with fresh LB (with the appropriate antibiotics and 0.2 % arabinose (w/v)) was inoculated with the cells at an OD_600_ of 0.01. The cells were grown aerobically in the Tecan Infinite F500 system (TECAN) at 37°C. OD_600_ and luminescence were recorded in 10 min intervals over the course of 16 h. Light units were normalized to OD_600_ and are thus expressed in relative lightunits (RLU). Each measurement was performed in triplicate.

To examine the regulatory role of rpoH, the growth temperature was adjusted to 20 °C and 25 °C during the measurement. The binding site for CAP was examined in LB with or without 20 mM glucose, thereby repressing or activating CyaA, respectively.

### Surface Plasmon Resonance Spectroscopy (SPR)

SPR spectroscopy and calibration free concentration (CFCA) assays were performed using a Biacore T200 device (GE Healthcare) and streptavidin-precoated Xantec SAD500-L carboxymethyl dextran sensor chips (XanTec Bioanalytics GmbH, Düsseldorf, Germany). All experiments were conducted at 25°C with HBS-EP+ buffer [10 mM HEPES pH 7.4, 150 mM NaCl, 3 mM EDTA and 0.05 % (v/v) detergent P20].

Before immobilizing the DNA fragments, the chips were equilibrated by three injections using 1 M NaCl/50 mM NaOH at a flow rate of 10 μl min^−1^. Then, 10 nM of the respective double-stranded biotinylated DNA fragment was injected using a contact time of 420 sec and a flow rate of 10 μl min^−1^. As a final wash step, 1 M NaCl/50 mM NaOH/50 % (v/v) isopropanol was injected. Approximately 250-350 RU of each respective DNA fragment were captured onto the respective flow cell. All interaction kinetics of FlrR with the respective DNA fragment were performed in HBS-EP buffer at 25°C at a flow rate of 30 μl min^−1^ in the presence and absence of 1mM 1 mM *N*-fructoselysine (ε-FrK). The proteins were diluted in HBS-EP buffer and passed over all flow cells in different concentrations (0.5 nM-125 nM) using a contact time of 180 s followed by a 300 sec dissociation time before the next cycle started. After each cycle the surface was regenerated by injection of 2.5 M NaCl for 30 sec at 60 μl min^−1^ flow rate followed by a second regeneration step by injection of 0.5 % (w/v) SDS for 30 sec at 60 μl min^−1^. All experiments were performed at 25°C. Sensorgrams were recorded using the Biacore T200 Control software 2.0 and analyzed with the Biacore T200 Evaluation software 2.0. The surface of flow cell 1 was not immobilized with DNA and used to obtain blank sensorgrams for subtraction of bulk refractive index background. The referenced sensorgrams were normalized to a baseline of 0. Peaks in the sensorgrams at the beginning and the end of the injection emerged from the runtime difference between the flow cells of each chip.

Calibration-free concentration analysis (CFCA) was performed using a 1 μM solution of purified FrlR, which was stepwise diluted 1:2, 1:5, 1:10, and 1:20. Each protein dilution was injected two-times, one at 5 μl min^−1^ as well as 100 μl min^−1^ flow rate. On the active flow cell DNA fragment including the P_*frlO*_-DNA was used for FlrR binding. CFCA basically relies on mass transport, which is a diffusion phenomenon that describes the movement of molecules between the solution and the surface. The CFCA therefore relies on the measurement of the observed binding rate during sample injection under partially or complete mass transport limited conditions. Overall, the initial binding rate (dR/dt) is measured at two different flow rates dependent on the diffusion constant of the protein. The diffusion coefficient of BceR-P was calculated using the Biacore diffusion constant calculator and converter webtool (https://www.biacore.com/lifesciences/Application_Support/online_support/Diffusion_Coefficient_Calculator/index.html), whereby a globular shape of the protein was assumed. The diffusion coefficient of FlrR was determined as D=1.02×10^−10^ m^2^/s. The initial rates of those dilutions that differed in a factor of at least 1.5 were considered for the calculation of the “active” concentration, which was determined as 5×10^−7^M (approximately 50 % of the total protein concentration) for FlrR. The “active” protein concentration was then used for calculation of the binding kinetic constants and steady-state affinity.

### Protein purification

His_6_-SUMO-tagged FraR from *S. enterica* (pET-SUMO-*fraR_S.en_*), FrlR from *E. coli* (pET-SUMO-*frlR_E.co_*) and *B. subtilis* (pET-SUMO-*frlR_B.su_*) were overproduced in *E.coli* BL21 (DE3) by addition of 1 mM IPTG to exponentially growing cells and subsequent cultivation at 18 °C o/n. Cells were lysed by sonication in the resepctive buffer (Table S4). The proteins were purified using Ni-nitrilotriacetic acid (Ni-NTA; Qiagen) according to the manufacturer’s instructions, using 20 mM imidazole for washing and 250 mM imidazole for elution. Subsequently, imidazole was removed by dialysis o/n at 4 °C in buffer 1. The His_6_-SUMO tag was cleaved by incubation with His_6_-Ulp1 (Starosta *et al.*, 2014) overnight. Subsequently, tag-free FrlR was collected from the flow through after metal chelate affinity chromatography.

Size exclusion chromatography was performed in the respective buffer (Table S4) using a Superdex 200 Increase 10/300-Gl column with a flow rate of 0.3 ml/min on an Äkta purifier (GE Healthcare). Four milligrams of protein was loaded in a volume of 0.4 ml (8,7 mg/ml). Eluting protein was detected at 280 nm. Fractions of 0.5 ml were collected.

### NMR experiments

All NMR spectra were acquired using an Bruker Avance III Bruker NMR spectrometer with a magnetic field strength corresponding to a proton Larmor frequency of 700 MHz Larmor frequency equipped with a room temperature triple resonance gradient probe head. NMR experiments were performed in a buffer containing 100 mM potassium phosphate, 300 mM NaCl, pH 6.5 at 298 K. Two dimensional 15N-HSQC titrations were performed with 200 μM FrlRUTRA domain in the absence and presence of 1.2 mM excess of ε-FrK. For measuring the T2 relaxation, one dimensional experiments were performed with 200 μM FrlR_UTRA_ in the presence of 1.2 mM J-FrK. 1D T2 experiments were performed using a 1-1 echo pulse sequence with a relaxation delay of varying between 0.004, 2.0 and 2.5 ms (Sklenář & Bax, 1987). ◻c was estimated using the equation 1/(5T2) where T2 is expressed in seconds (Kay et al., 1989, Barbato et al., 1992). Saturation transfer difference NMR experiments (Mayer & Meyer, 1999) were performed on 10 μM of FrlRUTRA + 1mM ε-FrK with irradiation at either 0.65 ppm (protein methyl region) or 8.5 ppm (protein amide region), far from ε-FrK signals to only saturate protein, with a relaxation delay of 5 s, which includes an effective saturation time of 4 s and an interscan delay of 1 s. For control experiments only 1mM ε-FrK was used.

### Fluorescence microscopy

The DNA-binding site of FrlR was examined by fluorescence microscopy of the strain BL21 (DE3) transformed with pUC19-PT7-O_*frlABCD*_-*sfGFP*-*frlR* or pUC19-PT7-*sfGFP*-*frlR*. The cells were cultivated overnight at 30 °C in LB medium supplemented with ampicillin. Expression of FrlR was induced or repressed by addition of 20 mM arabinose or 20 mM glucose, respectively. The overnight cultures were used to inoculate (OD_600_ of 0.1) fresh LB medium supplemented with ampicillin and arabinose or glucose. Cells were aerobically cultivated at 37◻°C and harvested by gentle centrifugation after 5 h. The pellet was washed twice and resuspended in PBS (OD_600_ 0.5). 2◻μl of the culture was spotted on 1 % (w/v) agarose pads, placed onto microscopic slides and covered with a coverslip. Subsequently, images were taken on a Leica DMi8 inverted microscope equipped with a Leica DFC365 FX camera (Wetzlar). An excitation wavelength of 484◻nm and a 535 nm emission filter with a 75-nm bandwidth was used for sfGFP fluorescence for 1 sec, gain 5, and 75 % intensity. Fluorescence intensity of at least 300 cells was measured and plotted using ImageJ.

### Thermal shift assay

5 μM of the Protein (FrlR (Eco)) was pipetted into 30 μl (2.5 μM) of DNA (cleaned up PCR product) on ice. 0.3 μl of a 1:10 dilution of SYPRO™ orange protein gel stain (ThermoFisher Scientific) was added to the mix. The mixture was transferred into a 96 well semi-skirted PCR plate. To ensure that the liquid was at the bottom the plate was centrifuged for 30 sec at 3000 rpm. Subsequently, the plate was inserted into iQ™5 Real-Time PCR Detection Systems (Bio-Rad). The 96 well plate was incubated at 8°C for 10 min. Every 30 seconds the temperature was increased by 0.5 °C until 90°C were reached. Fluorescence was recorded at a wavelength of 520 nm.

## Acknowledgements

We thank Ralph Krafczyk and Kirsten Jung for fruitful discussions.

## Funding

JL gratefully acknowledges financial support by the DFG research grant LA 3658/1-1. J.H. acknowledges support from the European Molecular Biology Laboratory (EMBL). PKAJ acknowledges EMBL and the EU Marie Curie Actions Cofund grant for an EIPOD fellowship.

## Conflict of interest

The authors declare no conflict of interest.

## Author contribution

Biochemical and genetic analyses on FrlR/FraR/NagR were conducted by N.G., F.K. and B.F.C.A.. NMR studies were performed by P.K.A.J and J.H. The corresponding proteins were produced and purified by B.F.C.A, N.G. and F.K.. M.H. and T.H. synthesized ARPs and analyzed ε/α-FrK turnover. J.L. designed the study. The manuscript was written by B.F.C.A, N.G. and F.K., P.K.A.J., J.H., M.H. and J.L..

## Data availability statement

All data generated or analysed during this study are included in this published article (and its supplementary information files).

