## Supplementary Tables S1-4 & Figures S1-S2 for "Transcriptional regulation of the *N*_ε_-fructoselysine metabolism in *Escherichia coli* by global and substrate-specific cues"

Supplement

**Supplementary table S1**: Primer used in this study

| Identifier | Oligonucleotide | Sequence (5' - 3') | Restriction site |
|---|---|---|---|
| | **Sequencing primers** | | |
| P1 | Seq33_fw | GGC GTC ACA CTT TGC TAT GC | |
| P2 | pBAD-HisA_rev | CAG TTC CCT ACT CTC GCA TG | |
| P3 | T7_prom | TAA TAC GAC TCA CTA TAG GG | |
| P4 | T7_term | TAT GCT AGT TAT TGC TCA G | |
| P5 | M13 rev | CAG GAA ACA GCT ATG ACC | |
| P6 | M13 uni | TGT AAA ACG ACG GCC A | |
| P7 | pUT18C_fw | CGG ATG TAC TGG AAA CGG TGC | |
| P8 | pUT18C_rev | CGG TGA AAA CCT CTG ACA CAT GC | |
| P9 | pKT25_fw | GCC ATT ATG CCG CAT CTG TCC A | |
| P10 | pKT25_rev | GCT GCA AGG CGA TTA AGT TGG G | |
| | **FrlABCD promotor contructs and thermoshift primer** | | |
| P11 | PspOMI_PfrlABCD-Rev | GCT ACC GGG CCC AGC GAT ACC TTT TTC TTG TCC AAT ACG | *PspOM*I |
| P12 | XbaI_PfrlABCD397_Fw | GCT CTA GAT CAT ATT GAA AAC CGT GAT AAG AAC TGC | *Xba*I |
| P13 | XbaI_PfrlABCD294_Fw | GCT CTA GAG TCT CTG ACA GAA ATC GGC TAA CA | *Xba*I |
| P14 | XbaI_PfrlABCD140_Fw | GCT CTA GAG ATC CTG TGC GAA ATT TTG TGA TCT TC | *Xba*I |
| P15 | XbaI_PfrlABCD108_Fw | GCT CTA GAC ATT ACA TAA CAT CAT ATG TTG TTA TAT TCA TCA T | *Xba*I |
| P16 | XbaI_PfrlABCD75_Fw | GCT CTA GAA TGC ATT GTC ATG TTA CCT TTT AAA TGA CTG | *Xba*I |
| P17 | XbaI_PfrlABCD50_Fw | TCT CTA GAT GAC TGC AAA CTC TCC CCT ACA | *Xba*I |
| P18 | PspOMI_PfrlABCD_-15_rev | GGT ACC GGG CCC TTG TCC AAT ACG TTG TAG GGG AG | *PspOM*I |
| P19 | PspOMI_PfrlABCD_-50_rev | GGT ACC GGG CCC TTT AAA AGG TAA CAT GAC AAT GCA TGA TG | *PspOM*I |
| P20 | PspOMI_PfrlABCD_-70_rev | GGT ACC GGG CCC TGC ATG ATG AAT ATA ACA ACA TAT GAT GTT A | *PspOM*I |
| P21 | EcoRI_PfrlABCD_294_Fw | GCT AGA ATT CGT CTC TGA CAG AAA TCG GCT AAC A | *EcoR*I |
| P22 | PfrlABCD_OL_sfGFP_OL_Rev | CTT TGC TCA TAG CGA TAC CTT TTT CTT GTC CAA TAC | |
| P23 | PfrlABCD_OL_sfGFP_OL_Fw | AGG TAT CGC TAT GAG CAA AGG AGA AGA ACT TTT CAC | |
| P24 | PspOMI_sfGFP_GS_His6_Rev | CTA GTG GGG CCC TTA GTG ATG GTG ATG GTG ATG CGA GCC TTT GTA GAG CTC ATC CAT GCC ATG | *PspOM*I |
| P25 | EcoR1_PT7_PfrlO_sRBS_Fw | GCC AGT GAA TTC TAA TAC GAC TCA CTA TAG GGT TGT CAT GTT ACC TTA TAA AAA AAT AAA TTA ATC GAA AGA TAG GCG CTA CGG TTA GGT AAA TGA GCA AAG GAG AAG AAC TTT TCA C | *EcoR*I |

| Identifier | Oligonucleotide | Sequence (5' - 3') | Restriction site |
|---|---|---|---|
| P26 | EcoR1_PT7_sRBS_Fw | GCC AGT GAA TTC TAA TAC GAC TCA CTA TAG GGA TAA AAA AAT AAA TTA ATC GAA AGA TAG GCG CTA CGG TTA GGT AAA TGA GCA AAG GAG AAG AAC TTT TCA C | *EcoR*I |
| | **Bacterial two hybrid constructs** | | |
| P27 | FrlR_XbaI_for | AC**T CTA GA**G ATG TCA GCT ACG GAC CG | *Xba*I |
| P28 | FrlR_KpnI_GS-His6_rev | AC**G GTA CC**C CAT GGT GAT GGT GAT GGT GGC TGC CAT GTC TTT TAT TAT CAA TAG TTA AGG T | *Knp*I |
| P29 | FrlR(Bsu)_XbaI_for | AC**T CTA GA**G TTA AAC AAC GGA AGT TCT ACA C | *Xba*I |
| P30 | FrlR(Bsu)_GS-His6_KpnI_rev | GT**G GTA CC**T CAT GGT GAT GGT GAT GGT GGC TTC CTG TGT AGG GGC TGT TG | *Kpn*I |
| P31 | FrlR-N_KpnI_GS-His6_rev | CC**G GTA CC**G TAT GGT GAT GGT GAT GGT GGC TTC CGC TTT GTA CAA AGG T | *Kpn*I |
| P32 | FrlR-C_XbaI_for | AC**T CTA GA**G AGC CAG AAA GTT GAA AAC G | *Xba*I |
| | **Protein overexpression** | | |
| P33 | SacI_sRBS-rpoH-Fw | CGC GAG CTC AAC ATC TGG AAA CAG ACT TAA CTA TAA GAG AGC CAA TTT TAT GAC TGA CAA AAT GCA AAG TTT AGC T | *Sac*I |
| P34 | XbaI_rpoH-Rev | CTC TAG ATT ACG CTT CAA TGG CAG CAC G | *Xba*I |
| P35 | FrlR(Eco)_Rev | TTA ATG TCT TTT ATT ATC AAT AGT TAA GGT GAT G | |
| P36 | FrlR(Eco)-Fw | TCA ACT ACG GAC CGC TAC TC | |
| P37 | FraR_Fw | AGT ATC GAG CAA CCC GAC AGT A | |
| P38 | FraR_Rev | TTA TAC GGT AAA CCG TAT TTT ATC GCC G | |
| P39 | FrlR(Eco)-S77_Fw | AGC CAG AAA GTT GAA AAC GCC CT | |
| P40 | FrlR(Bsu)-Fw | AGC TTA AAC AAC GGA AGT TCT ACA CCT TTA TAC | |
| P41 | FrlR(Bsu)-Rev | TTA TGT GTA GGG GCT GTT GAT GGT | |
| | **Surface plasmid resonance primer** | | |
| P42 | Bio-FrlO-Fw | [Btn]TGT TAT ATT CAT CAT GCA TTG TCA TGT TAC CTT TTA AAT GAC TGC AAA CT | |
| P43 | Bio-Random-Fw | [Btn]ACG GCC AGT GAA TTC TAA TAC GAC TCA CTA TAG GGA TAA AAA AAT AAA TT | |
| P44 | FrlO-Rev | AGT TTG CAG TCA TTT AAA AGG TAA CAT GAC AAT GCA TGA TGA ATA TAA CA | |
| P45 | Random-Rev | AAT TTA TTT TTT TAT CCC TAT AGT GAG TCG TAT TAG AAT TCA CTG GCC GT | |
| | **Reporterassay plasmids** | | |

| Identifier | Oligonucleotide | Sequence (5' - 3') | Restriction site |
|---|---|---|---|
| P46 | SacI-sRBS-FrlR_Fw | TTC GAG CTC CCA AGT TTT TAC TAC CCC CCC CAT ATA AGA TAC ATA ATA ACG AGG TAA ATA TGT CAG CTA CGG ACC GCT ACT C | *Sac*I |
| P47 | XbaI-sRBS-FrlR-GS-His6_Rev | CGA CTC TAG ATT AGT GAT GGT GAT GGT GAT GGC TGC CAT GTC TTT TAT TAT CAA TAG TTA AGG TGA TGC | *Xba*I |
| P48 | SacI_sRBS-FrlR(Bsu)-Fw | CGC GAG CTC TCC TAT TAC ACA AAA GTT CGT TAA GGA GGT TTT TTA TGT TAA ACA ACG GAA GTT CTA CAC CTT | *Sac*I |
| P49 | FrlR(Bsu)-GS_His6_Rev | CTC TAG ATT AGT GAT GGT GAT GGT GAT GGC TGC CTG TGT AGG GGC TGT TGA TGG TAA A | |
| P50 | XbaI-sRBS-FrlR-DNA-GS-His6_Rev | ACT CTA GAT TAG TGA TGG TGA TGG TGA TGG CTG CCG CTT TGT ACA AAG GT | *Xba*I |
| P51 | SacI-sRBS-FraR_ | TGC GAG CTC GCA GAC ATC TCT GCA TAT TGG GAT ATT TAA GGA GGT ATA TAT GAT CGA GCA ACC CGA CAG TA | *Sac*I |
| P52 | XbaI-GS-His6-FraR_Rev | CTC TAG ATT AGT GAT GGT GAT GGT GAT GGC TGC CTA CGG TAA ACC GTA TTT TAT CGC CG | *Xba*I |
| P53 | FrlR-N39A_Fw | CCT ACC GAA GCG GAG CTT TGT ACA | |
| P54 | FrlR-N39A_Rev | TGT ACA AAG CTC CGC TTC GGT AGG | |
| P55 | FrlR-R49A_Fw | AAC GTC AGC GCG ATT ACC ATT C | |
| P56 | FrlR-R49A_Rev | GAA TGG TAA TCG CGC TGA CGT T | |
| P57 | FrlR-I50A_Fw | GTC AGC CGC GCG ACC ATT CGC | |
| P58 | FrlR-I50A_Rev | GCG AAT GGT CGC GCG GCT GAC | |
| P59 | FrlR-R67A_Fw | GTA CTG ATC GCG TGG CAG GGA | |
| P60 | FrlR-R67A_Rev | TCC CTG CCA CGC GAT CAG TAC | |
| P61 | FrlR-G70A_Fw | CGC TGG CAG GCG AAA GGC A | |
| P62 | FrlR-G70A_Rev | TGC CTT TCG CCT GCC AGC G | |
| P63 | FrlR-K71A_Fw | TGG CAG GGA GCG GGC ACC TTT | |
| P64 | FrlR-K71A_Rev | AAA GGT GCC CGC TCC CTG CCA | |
| P65 | FrlR-T91A_Fw | GTG GTT TTG CGG ATT TTG GCG TC | |
| P66 | FrlR-T91A_Rev | GAC GCC AAA ATC CGC AAA ACC AC | |
| P67 | FrlR-R133A_Fw | CTC TGC GCG GTG ATG TAT CTC G | |
| P68 | FrlR-R133A_Rev | CGA GAT ACA TCA CCG CGC AGA G | |
| P69 | FrlR-S166A_Fw | GAA GGA AGC GCG ACC TAT CAG | |
| P70 | FrlR-S166A_Rev | CTG ATA GGT CGC GCT TCC TTC | |
| P71 | FrlR-Y168A_Fw | GCT CCA CCG CGC AGT TAT TTC | |
| P72 | FrlR-Y168A_Rev | GAA ATA ACT GCG CGG TGG AGC | |
| P73 | FrlR-D182A_Fw | GTG GTC AGC GCG AAA AAG ACC | |
| P74 | FrlR-D182A_Rev | GGT CTT TTT CGC GCT GAC CAC | |
| P75 | FrlR-N240A_Fw | TTA ACT ATT GAT GCG AAA AGA CAT GGC | |
| P76 | FrlR-N240A_Rev | GCC ATG TCT TTT CGC ATC AAT AGT TAA | |
| P77 | XbaI-frlD-(Eco)_Fw | ATG GAA TCT AGA TAA AAC CCT GGC GAC AAT CGG CG | *Xba*I |

| Identifier | Oligonucleotide | Sequence (5' - 3') | Restriction site |
|---|---|---|---|
| P78 | PtsI-frlD_(Eco)-GS-His6_rev | GAC GAC CTG CAG TTA GTG ATG GTG ATG GTG ATG GCT GCC CCA GGC ACC GTG GTA CTG AAT G | *Pst*I |
| P79 | XbaI-frlD-(Bsu)_Fw | ATG GAA TCT AGA TAA ATT GAT TGC GGT TGG AGA TAA TGT TGT AG | *Xba*I |
| P80 | PtsI-frlD-(Bsu)-GS-His6_Rev | GAC GAC CTG CAG TTA GTG ATG GTG ATG GTG ATG GCT GCC TAG TAT TCT CGT TTT TTC ACT GCT TCC CC | *Pst*I |
| P81 | XmaI-sRBS-nagR_Fw | GCC CGG GAC TGT AAC GAC CGG GGT CGT ATT TAA CTA AGA TCA GGT TTT TTA TGA ATA TCA ATA AAC AAT CGC CTA TTC C | *Xma*I |
| P82 | XbaI-nagR-GS-His6_Rev | CTC TAG ATT AGT GAT GGT GAT GGT GAT GGC TGC CTG AAA GAC GAT CCA TAT AGT GGA CAA AT | *Xba*I |

**Supplementary table S2:** Plasmids used in this study

| Plasmid | Feature/ Construction comments | Reference |
|---|---|---|
| Vector backbones | | |
| pBAD33 | Cam$^R$-cassette, p15A origin, AraC coding sequence, ara operator | (Guzman *et al.*, 1995) |
| pBAD24 | Amp$^R$-cassette, pBBR322 origin, AraC coding sequence, ara operator | (Guzman *et al.*, 1995) |
| pBBR1-MCS5 | Gm$^R$-cassette, pBBR broad host range origin of replication, mob region for conjugative transfer | (Kovach *et al.*, 1995) |
| pET-SUMO | Kan$^R$-cassette, pBR322 origin, small ubiquitin-like modifier (SUMO) | Invitrogen |
| pUC19 | Amp$^R$-cassette, pBR322 origin, lac promoter | (Yanisch-Perron *et al.*, 1985) |
| pKT25 | Kan$^R$-cassette, T25 fragment under transcriptional control of a *lac* promoter, MCS sequence at the 3' end (C-terminal) of T25 | EUROMEDEX |
| pKNT25 | Kan$^R$-cassette, T25 fragment under the transcriptional control of a *lac* promoter, MCS is at the 5’ end (N-terminal) of the fragment | EUROMEDEX |
| pUT18 | Amp$^R$-cassette, T18 fragment under the transcriptional control of a *lac* promoter, MCS is at the 5’ end (N-terminal) of the fragment | EUROMEDEX |
| pUT18C | Amp$^R$-cassette, T18 fragment under the transcriptional control of a *lac* promoter, MCS is at the 3’ end (C-terminal) of the fragment | EUROMEDEX |
| pBBR1-MCS5-TT-RBS-*lux* | Gm$^R$-cassette, Broad host range cloning vector; contains *luxCDABE* | (Gödeke *et al.*, 2011) |
| Bacterial two-hybrid analysis | | |
| pKT25-zip | Kan$^R$-cassette, control plasmid, leucine zipper of GCN4 genetically fused in frame to the T25 fragment (inserted within the *Kpn*I site of pKT25) | EUROMEDEX |

| Plasmid | Feature/ Construction comments | Reference |
|---|---|---|
| pUT18C-zip | $Amp^R$-cassette, control plasmid, leucine zipper of GCN4 genetically fused in frame to the T18 fragment (inserted between the *Kpn*I and the *EcoR*I site of pUT18C) | EUROMEDEX |
| pKT25 $FrlR_{Eco}$ | $Kan^R$-cassette, production vector, $FrlR_{Eco}$ fused to the 3' end of T25 / $frlR_{Eco}$ was amplified using P27 + P28 and cloned into pKT25 using the restriction enzymes as indicated by the primer name | this study |
| pKT25 FrlR-N | $Kan^R$-cassette, production vector, N-domain of $FrlR_{Eco}$ fused to the 3' end of T25 / *frlR*-N domain was amplified using P27 + P31 and cloned into pKT25 using the restriction enzymes as indicated by the primer name | this study |
| pKT25 FrlR-C | $Kan^R$-cassette, production vector, C-domain of $FrlR_{Eco}$ fused to the 3' end of T25 / *frlR*-C domain was amplified using P32 + P28 and cloned into pKT25 using the restriction enzymes as indicated by the primer name | this study |
| pKNT25 $FrlR_{Eco}$ | $Kan^R$-cassette, production vector, $FrlR_{Eco}$ fused to the 5' end of T25 / $frlR_{Eco}$ was amplified using P27 + P28 and cloned into pKNT25 using the restriction enzymes as indicated by the primer name | this study |
| pUT18C $FrlR_{Eco}$ | $Amp^R$-cassette, production vector, $FrlR_{Eco}$ fused to the 3' end of T25 / $frlR_{Eco}$ was amplified using P27 + P28 and cloned into pUT18C using the restriction enzymes as indicated by the primer name | this study |
| pUT18C FrlR-N | $Amp^R$-cassette, production vector, N-domain of $FrlR_{Eco}$ fused to the 3' end of T25 / *frlR*-N domain was amplified using P27 + P31 and cloned into pUT18C using the restriction enzymes as indicated by the primer name | this study |
| pUT18C FrlR-C | $Amp^R$-cassette, production vector, C-domain of $FrlR_{Eco}$ fused to the 3' end of T25 / *frlR*-C domain was amplified using P32 + P28 and cloned into pUT18C using the restriction enzymes as indicated by the primer name | this study |
| Reporter assays | | |
| pBAD33 $FrlR_{Eco}$ | $Cam^R$-cassette, production of *E. coli* FrlR used for reporter assays / $frlR_{Eco}$ was amplified with P46 + P47 and cloned into pBAD33 using the restriction enzymes as indicated by the primer name | this study |
| pBAD33 $FrlR_{Bsu}$ | $Cam^R$-cassette, production of *B. subtilis* FrlR used for reporter assays / $frlR_{Bsu}$ was amplified with P48 + P49 and cloned into pBAD33 using the restriction enzymes as indicated by the primer name | this study |
| pBAD33 rpoH (σ32) | $Cam^R$-cassette, production vector, arabinose inducible *raBAD* promoter ($P_{BAD}$) / *rpoH* was amplified with P33 + P34 and cloned into pBAD33 using the restriction enzymes as indicated by the primer name | this study |

| Plasmid | Feature/ Construction comments | Reference |
|---|---|---|
| pUC19 PT7 sfGFP $FrlR_{Eco}$ | $Amp^R$-cassette, production vector, sfGFP under the control of T7 polymerase promoter, *frlR*$_{Eco}$ under the control of an arabinose inducible promoter | this study |
| pUC19 PT7 PfrlO sfGFP $FrlR_{Eco}$ | $Amp^R$-cassette, production vector, sfGFP under the control of T7 polymerase promoter, *frlR*$_{Eco}$ under the control of an arabinose inducible promoter, $FrlR_{Eco}$ binding sequence upstream of the sfGFP | this study |
| pUC19 PT7 sfGFP | $Amp^R$-cassette, production vector, sfGFP under the control of T7 polymerase promoter / Insert was amplified with P23 + P24 and P26 + P22 and cloned into pUC19 using the restriction enzymes as indicated by the primer name | this study |
| pUC19 PT7 PfrlO sfGFP | $Amp^R$-cassette, production vector, sfGFP under the control of T7 polymerase promoter, $FrlR_{Eco}$ binding sequence upstream of the sfGFP / Insert was amplified with P23 + P24 and P25 + P22 and cloned into pUC19 using the restriction enzymes as indicated by the primer name | this study |
| pBBR MCS5 Lux Pfrl397 | Reporter construct containg the $P_{frlABCD}$ - promoter fusion -322/+75; $Gm^R$-cassette, Broad host range cloning vector; contains *luxCDABE* with upstream promoter sequence of the *frlABCDR* operon (397bp) / $P_{frlABCD}$ - promoter fusion was amplified with P11 + P12 and cloned into pBBR1-MCS5-TT-RBS-lux using the restriction enzymes as indicated by the primer name | this study |
| pBBR MCS5 Lux Pfrl294 | Reporter construct containg the $P_{frlABCD}$ - promoter fusion -219/+75; $Gm^R$-cassette, Broad host range cloning vector; contains *luxCDABE* with upstream promoter sequence of the *frlABCDR* operon (294bp) / $P_{frlABCD}$ - promoter fusion was amplified with P11 + P13 and cloned into pBBR1-MCS5-TT-RBS-lux using the restriction enzymes as indicated by the primer name | this study |
| pBBR MCS5 Lux Pfrl140 | Reporter construct containg the $P_{frlABCD}$ - promoter fusion -65/+75; $Gm^R$-cassette, Broad host range cloning vector; contains *luxCDABE* with upstream promoter sequence of the *frlABCDR* operon (140 bp) / P*frlABCD* - promoter fusion was amplified with P11 + P14 and cloned into pBBR1-MCS5-TT-RBS-lux using the restriction enzymes as indicated by the primer name | this study |
| pBBR MCS5 Lux Pfrl107 | Reporter construct containg the P*frlABCD* - promoter fusion -32/+75; $Gm^R$-cassette, Broad host range cloning vector; contains *luxCDABE* with upstream promoter sequence of the *frlABCDR* operon (108 bp) / P*frlABCD* - promoter fusion was amplified with P11 + P15 and cloned into pBBR1-MCS5-TT-RBS-lux using the restriction enzymes as indicated by the primer name | this study |

| Plasmid | Feature/ Construction comments | Reference |
|---|---|---|
| pBBR MCS5 Lux Pfrl15-294 | Reporter construct containg the $P_{frlABCD}$ - promoter fusion -219/+60; $Gm^R$-cassette, Broad host range cloning vector; contains *luxCDABE* with upstream promoter sequence of the *frlABCD* operon (15-294 bp) / $P_{frlABCD}$ - promoter fusion was amplified with P21 + P18 and cloned into pBBR1-MCS5-TT-RBS-lux using the restriction enzymes as indicated by the primer name | this study |
| pBBR MCS5 Lux Pfrl50-294 | Reporter construct containg the $P_{frlABCD}$ - promoter fusion -219/+25; $Gm^R$-cassette, Broad host range cloning vector; contains *luxCDABE* with upstream promoter sequence of the *frlABCD* operon (50-294 bp) / $P_{frlABCD}$ - promoter fusion was amplified with P21 + P19 and cloned into pBBR1-MCS5-TT-RBS-lux using the restriction enzymes as indicated by the primer name | this study |
| pBAD33-frlR-DNA$_{Eco}$ | $Cam^R$-cassette, Production of *E. coli* FrlR DNA binding domain used for reporter assays / *frlR*-DNA$_{Eco}$ domain was amplified with P46 + P50 and cloned into pBAD33 using the restriction enzymes as indicated by the primer name | this study |
| pBAD33-fraR$_{Sen}$ | $Cam^R$-cassette, Production of *S. Typhimurium* FraR used for reporter assays / *fraR*$_{Sen}$ domain was amplified with P51 + P52 and cloned into pBAD33 (*Sac*I/*Xba*I) | this study |
| pBAD33-nagR$_{Bsu}$ | $Cam^R$-cassette, Production of *B. subtilis* NagR used for reporter assays / *nagR*$_{Bsu}$ domain was amplified with P81 + P82 and cloned into pBAD33using the restriction enzymes as indicated by the primer name | this study |
| pBAD33-frlR-N39A$_{Eco}$ | $Cam^R$-cassette, Production of *E. coli* mutant frlR- N39A used for reporter assays / frlR$_{Eco}$ variant was amplified with P46+P54 and P53+P47 and cloned into pBAD33 using the restriction enzymes as indicated by the primer name | this study |
| pBAD33-frlR-R49A$_{Eco}$ | $Cam^R$-cassette, Production of *E. coli* mutant FrlR- R49A used for reporter assays / *frlR*$_{Eco}$ variant was amplified with P46+P56 and P55+P47 and cloned into pBAD33 using the restriction enzymes as indicated by the primer name | this study |
| pBAD33-frlR-I50A$_{Eco}$ | CamR-cassette, Production of *E. coli* mutant FrlR- I50A used for reporter assays / *frlR*$_{Eco}$ variant was amplified with P46+P58 and P57+P47 and cloned into pBAD33 using the restriction enzymes as indicated by the primer name | this study |
| pBAD33-frlR-R67A$_{Eco}$ | $Cam^R$-cassette, Production of *E. coli* mutant FrlR- R67A used for reporter assays / *frlR*$_{Eco}$ variant was amplified with P46+P60 and P59+P47 and cloned into pBAD33 using the restriction enzymes as indicated by the primer name | this study |

| Plasmid | Feature/ Construction comments | Reference |
|---|---|---|
| pBAD33-frlR-G70A$_{Eco}$ | Cam$^R$-cassette, Production of *E. coli* mutant FrlR- G70A used for reporter assays / *frlR*$_{Eco}$ variant was amplified with P46+P62 and P61+P47 and cloned into pBAD33 using the restriction enzymes as indicated by the primer name | this study |
| pBAD33-frlR-K71A$_{Eco}$ | Cam$^R$-cassette, Production of *E. coli* mutant FrlR- K71A used for reporter assays / *frlR*$_{Eco}$ variant was amplified with P46+P64 and P63+P47 and cloned into pBAD33 using the restriction enzymes as indicated by the primer name | this study |
| pBAD33-frlR-T91A$_{Eco}$ | Cam$^R$-cassette, Production of *E. coli* mutant FrlR- T91A used for reporter assays / *frlR*$_{Eco}$ variant was amplified with P46+P66 and P65+P47 and cloned into pBAD33 using the restriction enzymes as indicated by the primer name | this study |
| pBAD33-frlR-R133A$_{Eco}$ | Cam$^R$-cassette, Production of *E. coli* mutant FrlR- R133A used for reporter assays / *frlR*$_{Eco}$ variant was amplified with P46+P68 and P67+P47 and cloned into pBAD33 using the restriction enzymes as indicated by the primer name | this study |
| pBAD33-frlR-S166A$_{Eco}$ | Cam$^R$-cassette, Production of *E. coli* mutant FrlR- S166A used for reporter assays / *frlR*$_{Eco}$ variant was amplified with P46+P70 and P69+P47 and cloned into pBAD33 using the restriction enzymes as indicated by the primer name | this study |
| pBAD33-frlR-Y168A$_{Eco}$ | Cam$^R$-cassette, Production of *E. coli* mutant FrlR- Y168A used for reporter assays / *frlR*$_{Eco}$ variant was amplified with P46+P72 and P71+P47 and cloned into pBAD33 using the restriction enzymes as indicated by the primer name | this study |
| pBAD33-frlR-D182A$_{Eco}$ | Cam$^R$-cassette, Production of *E. coli* mutant FrlR- D182A used for reporter assays / *frlR*$_{Eco}$ variant was amplified with P46+P74 and P73+P47 and cloned into pBAD33 using the restriction enzymes as indicated by the primer name | this study |
| pBAD33-frlR-N240A$_{Eco}$ | Cam$^R$-cassette, Production of *E. coli* mutant frlR- N240A used for reporter assays / *frlR*$_{Eco}$ variant was amplified with P46+P76 and P75+P47 and cloned into pBAD33 using the restriction enzymes as indicated by the primer name | this study |
| pBAD24-frlD$_{Eco}$ | Amp$^R$-cassette, Production of *E. coli* FrlD used for reporter assays and BW25113 Δ*frlD* complementation / frlD$_{Eco}$ domain amplified with P77+P78 and cloned into pBAD24 using the restriction enzymes as indicated by the primer name | this study |
| pBAD24-frlD$_{Bsu}$ | Amp$^R$-cassette, Production of *B. subtilis* FrlD used for reporter assays and BW25113 Δ*frlD* complementation / *frlD*$_{Bsu}$ domain amplified withP79+P80 and cloned into pBAD24 using the restriction enzymes as indicated by the primer name | this study |

| Plasmid | Feature/ Construction comments | Reference |
|---|---|---|
| Protein production | | |
| pET-SUMO-frlR$_{Eco}$ | Kan$^R$-cassette, Production of *E. coli* FrlR used for protein purification, *frlR$_{Eco}$* domain was amplified using P35 + P36 using the restriction enzymes as indicated by the primer name | this study |
| pET-SUMO-frlR$_{Bsu}$ | Kan$^R$-cassette, Production of *B. subtilis* FrlR used for protein purification, frlR$_{Eco}$ domain was amplified using P40 + P41 using the restriction enzymes as indicated by the primer name | this study |
| pET-SUMO-fraR$_{Sen}$ | Kan$^R$-cassette, Production of *S. enterica* FraR used for protein purification and characterization / fraR$_{Sen}$ amplified with P37+P38 and cloned into pET-SUMO using the restriction enzymes as indicated by the primer name | this study |
| pET-SUMO-frlR-UTRA$_{Eco}$ | Kan$^R$-cassette, Production of *E. coli* FrlR-UTRA domain used for protein purification and characterization / *frlR*-UTRA domain$_{Eco}$ amplified with P39+P35 and cloned into pET-SUMO using the restriction enzymes as indicated by the primer name | this study |

**Supplementary table S3:** Strains used in this study

| Strain | Feature/Construction comments | Source |
|---|---|---|
| *E. coli* DH5αλpir | F⁻ ϕ80*lac*ZΔM15 Δ(*lac*ZYA-*arg*F)U169 *rec*A1 *end*A1 *hsd*R17($r_K^-$, $m_K^+$) *pho*A *sup*E44 λ⁻ *thi*-1 *gyr*A96 *rel*A1, *uidA*::*pir*+ | (Macinga *et al.*, 1995) |
| *E. coli* BL21(DE3) | F⁻ *omp*T *hsd*S$_B$ ($r_B^-$, $m_B^-$) *gal dcm* (DE3) | (Studier & Moffatt, 1986) |
| *E. coli* BW25113 | F-, Δ*(araD-araB)*567, Δ*lacZ4787*(::rrnB-3), *λ*$^-$, Δ*frlD781::kan*, *rph-1*, Δ*(rhaD-rhaB)568*, *hsdR514* | (Datsenko & Wanner, 2000) |
| *E. coli* BTH101 | F-, *cya-99, araD139, galE15, galK16, rpsL1 (Str$^r$), hsdR2, mcrA1, mcrB1* | Euromedex |
| JW5698 | BW25113; Δ*frlR*782::kan | (Baba *et al.*, 2006) |
| JW3337 | BW25113; Δ*frlD*781::kan | (Baba *et al.*, 2006) |
| JW3778 | BW25113; Δ*cyaA*751::kan | (Baba *et al.*, 2006) |

**Supplementary table S4:** Buffer used in this study

| Buffer<br>pH | Concentration | Compound |
|---|---|---|
| **Buffer FrlR** | | |
| pH 7.5 | 300 mM<br>100 mM<br>5 mM<br>1 mM<br>10% (v/v) | NaCl<br>NaPi pH 7.5<br>DTT<br>PMSF<br>Glycerol |
| **Buffer FraR** | | |
| pH 8.2 | 0.1 M<br>0.3 M<br>10% (v/v) | NaPi<br>NaCl<br>Glycerol |
| **Buffer SpeC** | | |
| pH 7.6 | 50 mM<br>300 mM<br>10% (v/v)<br>1 mM | Tris<br>NaCl<br>Glycerol<br>DTT |

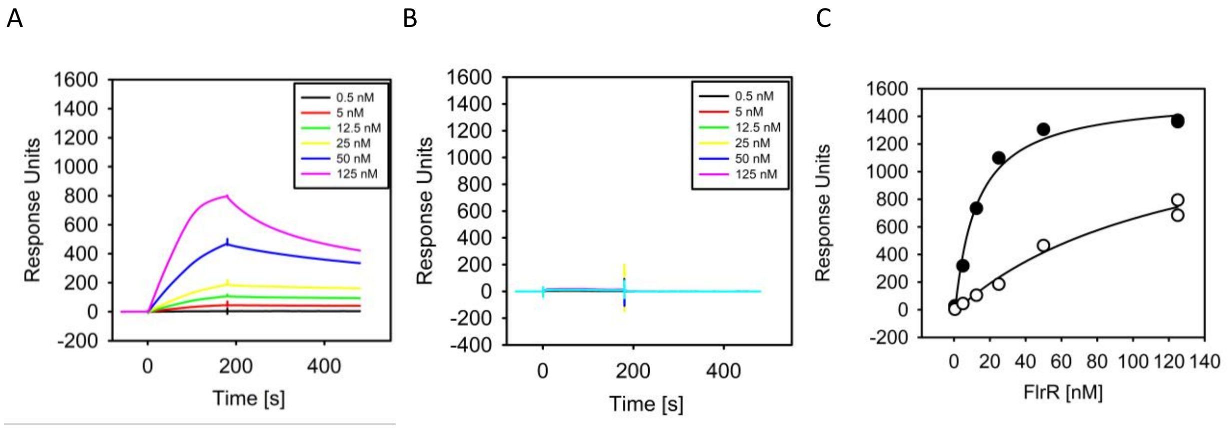


**Fig. S1:** *In vitro* DNA binding of FrlR to $P_{frlO}$ analyzed by surface plasmon resonance spectroscopy (SPR) in the presence of 1 mM $N_{\varepsilon}$-fructoselysine (ε-FrK): The biotin-labeled DNA fragment $P_{flrO}$ (A), and the control fragment without the FlrR binding motif (B) were captured on a streptavidin-coated sensor chip, and purified FlrR was passed over the chip at a flow rate of 30 µl/min and temperature of 25°C [concentrations of 0.5 nM, 5 nM, 12.5 nM, 25 nM, 50 nM, 125 nM), using a contact (association) time of 180 sec, followed by a 300-sec dissociation phase. The increase in RU correlates with an increasing FlrR concentration. (C): Steady state binding of FlrR to $P_{flrO}$ in the presence (white circles) and absence (black circles) of ε-FrK.

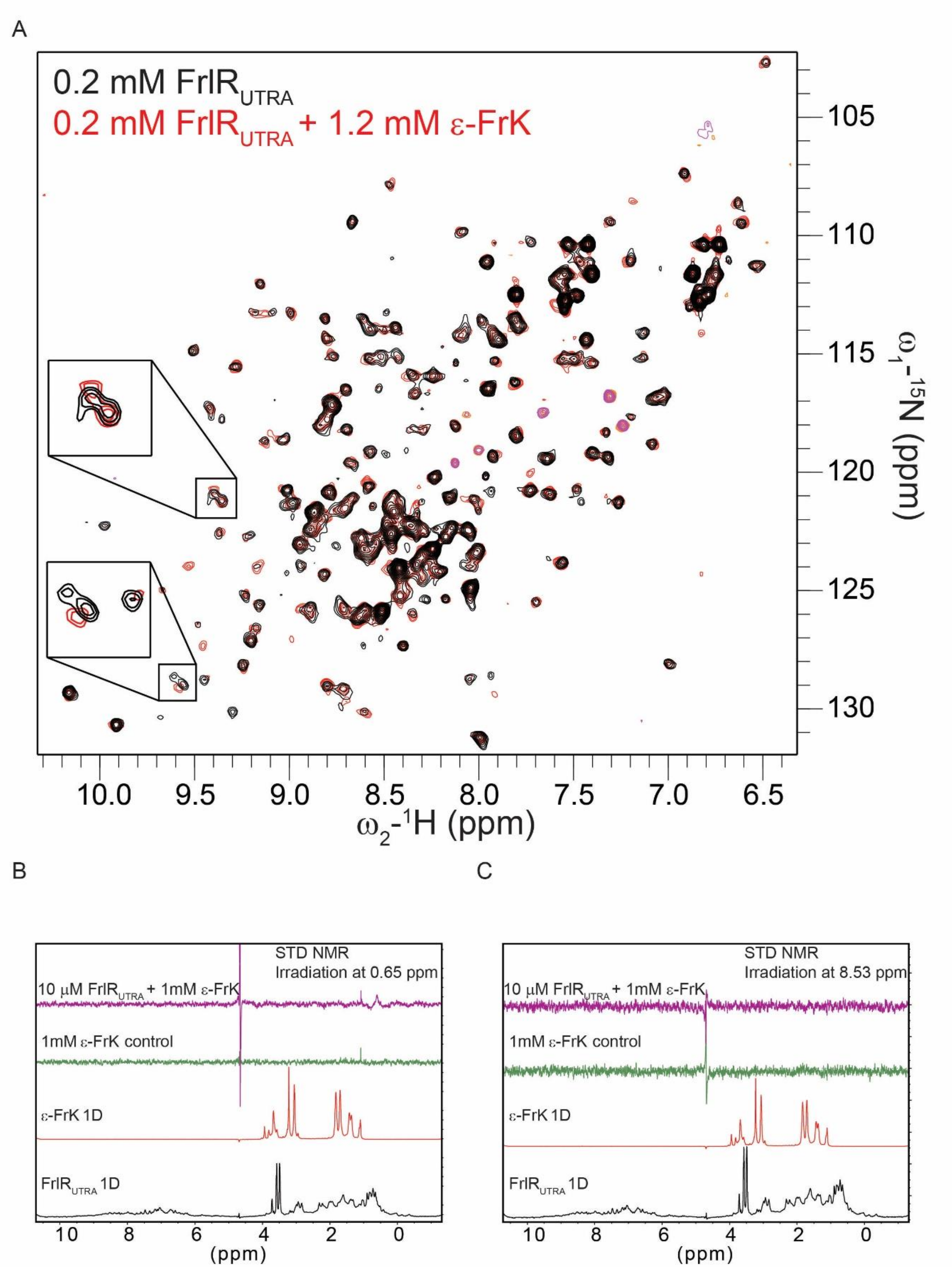


**Fig. S2: NMR investigation of FrlR and FrlR$_{UTRA}$ and binding to ε-FrK.** A) FrlR$_{UTRA}$ $^{1}$H-$^{15}$N HSQC spectrum in absence (black) and 6-fold excess of ε-FrK (red). There are only minor chemical shift perturbations upon titration suggesting a weak interaction of ε-FrK with FrlR$_{UTRA}$. B and C) Saturation transfer difference (STD)-NMR experiments with 10 µM FrlR$_{UTRA}$ and 1mM ε-FrK , irradiating with Gaussian pulses at 0.65 ppm (D) and 8.53 ppm (E). No STD signals could be detected, confirming that the affinity of ε-FrK for FrlR$_{UTRA}$ is weak and lower then $10^{-3}$ M.
